# Nesting flight statistics for wind turbine planning: a MoveApps workflow

**DOI:** 10.1101/2023.01.27.525824

**Authors:** Andrea Kölzsch, Johannes Gal

## Abstract

As green, renewable energy is increasing by the installation of more and more wind turbines, the assessment of their impact on protected species has to be improved by more automatized, data-driven risk analyses.

We have developed as set of two workflows to extract simple parameters for collision risk models from GPS tracks of sensitive bird species during nesting. The workflows have been integrated into the free MoveApps platform and are available there. The analysis code of all components of the workflows is openly available on GitHub, and improvement and adaption to other, similar requirements is encouraged.

With three example data sets of white storks (WS), red kites (RK) and marsh harriers (MH), we illustrate how the workflows are used. The first workflow identifies nesting sites and time of nesting from the GPS tracks, the second workflow calculates flight speeds, flight duration, flight height and distance from the nest. Estimated flight speeds show low within species variability, with averages of 4.1 m/s (MH), 6.9 m/s (RK) and 10.9 m/s (WS). Extracted times of nesting are widely spread through the season and flight height and distance to the nest when in flight show large differences between individuals and years. Similar to the central evaluation distances around the nest required by national legislation, the 50% in-flight usage thresholds are 700 m (MH), 1100 m (RK) and 1400 m (WS). Flight height during nesting is rather low, on average 16 m (MH), 75 m (RK) and 193 m (WS) above the ground.

These values can help to estimate collision risk with wind turbines for large birds in Central Europe during their nesting period. Finally, the developed MoveApps workflows (an possible adaptions) can be used to extract required parameters from tracking studies of any other vulnerable species or populations in an unbiased, automated manner to improve wind turbine placement in relation to nesting sites.

## Introduction

To lower emissions that presently cause and increase climate change, the use of renewable energy must increase (Schleussner et al. 2016). One of the most effective green energy technologies are wind turbines (Kumar et al. 2016), which are constructed in great numbers on the land or off-shore. Wind turbines have, however, been reported to impact bird and bat species (Saidur et al. 2011; Smith and Dwyer 2016), mainly by increasing mortality by collision strikes (Drewitt and Langston 2008; Thaxter et al. 2017). Significantly increased mortality of wildlife is contrary to e.g. the German Federal Nature Conservation Act (Federal Law Gazette 2009) and various conservation efforts (Kuvlesky Jr. et al. 2007). Therefore, the planning of new wind farms and the evaluation of suitable placement of wind turbines must include an assessment of the mortality risk of sensitive species (Smallwood 2007).

Reproduction is one of the most important life stages for animals and, as they spend most time in areas close to their nests then, they are especially vulnerable to any disturbances in their breeding sites. Especially large birds can be impacted by the turning rotors of wind turbines close to their nests. Thus, legislation in e.g. Germany has focussed on the evaluation of the impact of wind turbines on large nesting birds, against the background of the distance between the nests and the potential wind turbine site. Previous approaches provide recommended distances between planned wind turbines and nesting sites (e.g. by the Working Group of Ornithological Observatories (Länderarbeitsgemeinschaft der Vogelschutzwarten 2014) and in § 45b of the Germany Federal Nature Conservation Act (Federal Law Gazette 2009)). However, in the past years, more complex collision risk models have been developed to improve impact assessment and generalise and alleviate procedures (Band 2012; Masden 2015; BDEW Bundesverband der Energie-und Wasserwirtschaft e.V. 2021). In order to deliver satisfactory results, the models require as input ascertained and generalized flight parameters of the species of concern.

Many GPS tracks have been collected by a large number of scientists, wildlife managers and conservation groups worldwide and uploaded and archived in online databases like Movebank (Kays et al. 2022). For some species, the data have been analysed in relation to collision risk (Schaub et al. 2020; Pfeiffer and Meyburg 2022), wind turbine avoidance behaviour (Chamberlain et al. 2006; Watson et al. 2018; Santos et al. 2022) and habitat loss (Marques et al. 2020). However, much available tracking data has not been analysed for wind turbine collision risk applications. With access to tracking databases, it is possible to assess the availability and complexity of the available data sets, to contact data owners and with their agreement use the tracks for additional analyses, like the extraction of generalised parameters as input for collision risk models.

Here, we present two workflows that allow easy extraction of simple, but relevant statistics from GPS tracking data and their results for three example data sets (White stork *Ciconia ciconia*, Red kite *Milvus milvus* and Marsh harrier *Circus aeruginosus*). The workflows are implemented in the open-source MoveApps platform, www.moveapps.org (Kölzsch et al. 2022) that allows seamless analysis of shared data from Movebank. The first workflow extracts nesting sites and nesting times from the tracks, so that only movement characteristics during this most important time period in the life of the birds is considered. The second workflow estimates flight speed, the daily proportion of time in flight, the proportion of flight in different heights and the proportions of presence when in flight at certain distances around the nest. These workflows may be used by anybody with access to tracking data and some background knowledge of the tracked species, and the extracted parameters can be used as input for collision risk models.

## Methods

### Data selection

Before being able to properly analyse any tracking data, it is important to get an overview of the data collection process, tag settings (e.g. resolution, gaps) and the species’ ecology. Parameter settings for nest detection and flight speed were defined in discussion with data owners (see below). We estimate that a minimum regular resolution of about 1 location per 20 minutes is required for sensible estimations of flight statistics, as flight durations are short during the nesting periods (own observations). Gaps in data collection are possible to account for, but limit the significance of the results. For all statistics below, it is crucial to use instantaneous GPS ground speed as transmitted by the GPS chip of the tags, because they are more accurate and characterise the behaviour of the animal at the time of location measurement.

Here, we analyse GPS data sets of white storks, red kites and marsh harriers, which are listed in Appendix 1 of the Federal Nature Conservation Act of Germany (Federal Law Gazette 2009) and considered to be sensitive towards the use of wind energy regarding collisions. All tracks of the selected data are stored on Movebank: (1) in the public study “Life Track White Stork SW Germany”, with tracks in Southern Germany in 2013-2022 (GPS resolution depending on battery charge 1 location per 20 minutes or 1 location per 5 minutes; no data collection at night), (2) in the non-public study “Red Kite MPI-AB Baden-Wuerttemberg”, with tracks in South-West Germany in 2018-2022 (GPS resolution depending on battery charge 1 location per 5 minutes or 1 location per 1 minute; no data collection at night) and (3) in a set of three public studies “H_GRONINGEN - Western marsh harriers (Circus aeruginosus, Accipitridae) breeding in Groningen (the Netherlands)”, “MH_WATERLAND - Western marsh harriers (Circus aeruginosus, Accipitridae) breeding near the Belgium-Netherlands border” and “BOP_RODENT - Rodent specialized birds of prey (Circus, Asio, Buteo) in Flanders (Belgium)”, with tracks in Belgium and the Netherlands in 2012-2018. GPS resolution depends on battery charge at time of collection and ranges between 1 location per 30, 15 and 5 minutes. Partly, locations were regularly collected also at night, but for some tracks one location per night (local midnight) was available. Whenever location bursts had been collected, we retained only the first value for our analyses.

## 1. Nest Location and Nesting Duration workflow

As a first step, nesting sites were extracted from the GPS tracks using the “Nest Location and Nesting Duration” workflow on MoveApps (Kölzsch and Flack 2022; see Appendix 1)). It starts with loading data from Movebank. As the nest detection algorithm in the final App of this workflow (see below) is based on many pairwise distance calculations that need much memory, we subsample the data for this analysis to 1 location per hour. Furthermore, only full breeding seasons are of interest for this analysis, therefore only tracks of animals with at least data of one complete breeding season are selected. In the third preparatory step, each track is split into single years/seasons using the Filter by Season App. All further calculations are done on those single year tracks.

The Nest Cluster Detection App is the centre of this workflow and is based on the R package nestR (Picardi et al. 2020). It needs a number of input parameters (Settings) regarding mainly the nesting duration, percentage of nest attendance, revisits and data quality, that we have evaluated together with species experts and iteratively adapted. Many failed breeding attempts are excluded. The method is conservative and robust to inaccuracies of those parameters such that nests would not be detected if they deviated from best values, leading to the respective track(s) not being included in the calculations of the second workflow (below). Breeding seasons are pre-defined to lie in the interval of 1 March – 31 August. For details about the algorithm see also the documentation of the underlying R package (Picardi et al. 2020). The nest_table.csv that is created by the Nest Cluster Detection App is the main output of this workflow, and is also used as input for the second workflow below. The table should be evaluated manually before further use.

## 2. Nest flight statistics workflow

Based on the detected nesting sites and time intervals of nest use, in the second MoveApps workflow (Kölzsch and Gal 2022, see details in Appendix 1) we calculate parameters of flight and nest use behaviour. The workflow starts with downloading data from Movebank in full resolution and splitting it into yearly nesting time tracks. Then, with help of the nest_table.csv, of each track only the individual-specific nesting intervals are extracted, ensuring that all calculations relate to the time that the birds were nesting. Note that it is also possible to manually prepare a nest table file from known nest sites and use it in this second workflow (see Appendix 1 for details).

First, flight speeds are estimated by fitting a bimodal to the distribution of GPS ground speed of each individual nesting track using the R package multimode. We use the turning point between the two peaks (antimode) as threshold between non-flight and flight locations and the second peak as individual average flight speed. Averages of these two individual values are calculated for each species and used in consecutive analyses.

Second, the duration that a bird was in flight every day during the nesting season indicates how often it might come close to nearby wind turbines. Therefore, our workflow calculates the proportion of locations with GPS ground speed above the average antimode (i.e. flight) per day for each individual and day, and determines averages. As the proportion of locations might be biased by unregular data collection, we also calculate daily cumulative flight durations, based on each location’s time lag to the next location, assuming that the same behaviour (flight/no flight) was retained for that duration. Before calculating these cumulative flight durations, we take gaps into account by adapting location time lags that are larger than 30 min to the median resolution of the respective data set. As a consequence, however, the daily proportions of flight behaviour are based on the total daily tracking duration (time that tags collected regular locations without e.g. night gap) instead of 24 hours.

Third, depending on the turbine height, flight height is important to estimate how often the birds fly in the dangerous zone of the turning blades. GPS tags collect, depending on the model and company, height above mean sea level or height above the earth ellipsoid. To convert this into height above ground, one has to adapt the height variable with ground heights. These are provided globally by Digital Elevation Models. Using the R package elevatr, the workflow appends ground elevation to each flight location (GPS ground speed above antimode) and calculates ground height from the available GPS height measurement. In addition to average flight heights, aggregated values of proportional use of flight height are provided, always cumulative between the selected height and the ground.

If the tags collected height above mean sea level (often called height above msl), the adapted value is relatively accurate. However, if height above ellipsoid is provided (as is the case here for the white stork data set), the inaccuracy is larger, in the range of 50-100 m. If higher accuracies are necessary, a transformation is advised. As the transformation between height above msl and height above ellipsoid is complex and differs strongly by region on the globe, we have so far not included this transformation in the workflow. It is planned for a future version under the use of the Earth Gravitational Model (https://earth-info.nga.mil/index.php?dir=wgs84&action=wgs84#tab_egm2008). The white stork data set that is analysed here contains height above ellipsoid only. For analyses in Germany (where all white stork tracks are measured), mean sea level is between 30-40 m above the ellipsoid (i.e. geoid height, see https://www.unavco.org/software/geodetic-utilities/geoid-height-calculator/geoid-height-calculator.html), so that to adapted ellipsoid height we add 30 m for the true ground height.

Fourth, to evaluate how far from a nest wind turbines can be erected, proportional use of the area around the nest site must be evaluated. This parameter has been provided as evaluation area radii in the German Federal Nature Conservation Act (Federal Law Gazette 2009) and estimated for the red kite by different studies (Mammen et al. 2013; Pfeiffer and Meyburg 2022), so we can compare them with our outcomes. Our workflow calculates the proportion of area used in different distances around the nest cumulatively for a set of circular distances.

## Results

### MoveApps workflows

The two developed MoveApps workflows – (1) Nest Location and Nesting Duration and (2) Nest Flight Statistics – have been publicly shared on MoveApps, so that anybody with tracking data of sufficient quality can use them and reproduce our analyses. Furthermore, the code of all Apps that make up the workflows is available openly on GitHub. Detailed instructions how to use the workflows in MoveApps are provided in Appendix 1. Both workflows have run times of between 20-60 minutes, which makes the use of a serverless cloud computing system like MoveApps highly valuable.

### Nest location and nesting duration

All data analysed in the three example use cases has been collected in Central Europe (Germany, Netherlands, Belgium). Our selected data sets consist of 28 tracks of 8 individual white storks, 25 tracks of 10 individual red kits and 12 tracks of 5 individual marsh harriers (Appendix 2-4).

The three species differ in breeding durations as well as nest attendance and revisit parameters (Table 1). Many of the detected nests were evaluated in the field and confirmed as known nest sites, especially for the white stork data. It is notable that nesting times varied strongly between individuals of all species (Appendix 2-4; Table 2), spreading across (almost) the complete five months of breeding season.

**Table 1.** Settings of the Nest Detection App (Picardi et al. 2020) for the three considered species.

|  | **White stork** | **Red kite** | **Marsh harrier** |
| --- | --- | --- | --- |
| **Duration of a nesting cycle (days)** | 150 | 85 | 70 |
| **Size of the buffer to compute location revisitation (m)** | 40 | 50 | 100 |
| **Minimum number of points within a buffer** | 5 | 5 | 5 |
| **Minimum number of fixes for a day to be retained of no recorded nest visit** | 5 | 5 | 3 |
| **Minimum number of consecutive days visited** | 80 | 35 | 30 |
| **Minimum percentage of fixes at a location on the day with maximum attendance** | 75 | 50 | 50 |
| **Minimum percentage of day spent at a location between first and last visit** | 1 | 1 | 5 |
| **Discard overlapping** | TRUE | TRUE | TRUE |

**Table 2.** Nesting and flight statistics of three example species. For plain statistics, average ± standard deviation is provided. See also Appendix 2-4 for individual values of the given properties.

|  | **White stork** | **Red kite** | **Marsh harrier** |
| --- | --- | --- | --- |
| **Start of breeding** | 01 March | 3 March | 31 March |
| **End of breeding** | 31 August | 31 August | 16 August |
| **Breeding duration** | 144.8 days (4.8 months) | 81.4 days (2.7 months) | 66.4 days (2.2 months) |
| **Flight speed** | 10.9 ($\pm 0.5$) m/s | 6.9 ($\pm 0.6$) m/s | 4.1 ($\pm 0.7$) m/s |
| **Flight speed threshold** | 4.6 ($\pm 0.9$) m/s | 2.3 ($\pm 0.5$) m/s | 2.3 ($\pm 0.7$) m/s |
| **Daily flight duration** | 0.75 ($\pm 0.27$) hours | 3.54 ($\pm 1.73$) hours | 5.4 ($\pm 0.7$) hours |
| **Daily tracking duration** | 17.1 ($\pm 1.8$) hours | 12.5 ($\pm 5.1$) hours | 16.2 ($\pm 5.0$) hours |
| **Flight height above ground** | 162.6 + 30 = 192.6 ($\pm 545.3$) m | 75.0 ($\pm 110.6$) m | 16.2 ($\pm 100.7$) m |
| **Individual flight height above ground (median = 50%)** | Between 105 and 130m | Between 50 and 75m | below 25m |
| **Distance to nest** | 3083 ($\pm 6820$) m | 1696 ($\pm 3564$) m | 1088 ($\pm 1841$) m |
| **Distance to nest (median = 50%)** | 1400 m | 1100 m | 700 m |

### Nest flight statistics

Flight speeds are very clearly discernible from GPS ground speed of each individual. The average values differ strongly between species (10.9 m/s for white stork, 6.9 m/s for red kite and 4.1 m/s for marsh harrier; Table 2), but have low standard deviation between individuals of the same species (0.5 – 0.7 m/s; Appendix 2-4). The same is true for flight speed thresholds. Thus, we are very confident to base the following analyses on these extracted values.

Daily flight durations differ strongly between white stork (45 min) and the other two species (3.5-5.4 hours; Appendix 2-4; Table 2), likely due to their different foraging modes: white storks feed on the ground, red kites and marsh harriers hunt in flight. Standard deviations of these values are rather high, which might be due to local conditions as well as differences in tag performance causing different daily tracking durations.

For white stork and red kite tracks, no data has been collected at the dark hours of the night, so daily tracking durations are less than 24 hours. It is expected that the birds do not move much and do not fly during this dark time, but it cannot be concluded from the data. For several species, unexpected flight behaviour during night has been recorded in the past, especially during nights with full moon. Therefore, our results are based on daily tracking durations instead of the 24 hours per day.

Flight height above the ground differs between all species, white storks flying highest (192.6 m; Table 2), which can be attributed to their ability and need of thermalling flight to save energy. Flight heights seem to be, however, rather variable (high standard deviations; Appendix 2-4; Table 2), depending on habitat, weather conditions and GPS inaccuracies. The distribution shape of flight heights is similar between species, with high usage of lower heights and decreasing usage of higher layers (Figure 1). The skew toward lower heights is most extreme for the marsh harrier that flies mostly below 25 m.

**Figure 1.**
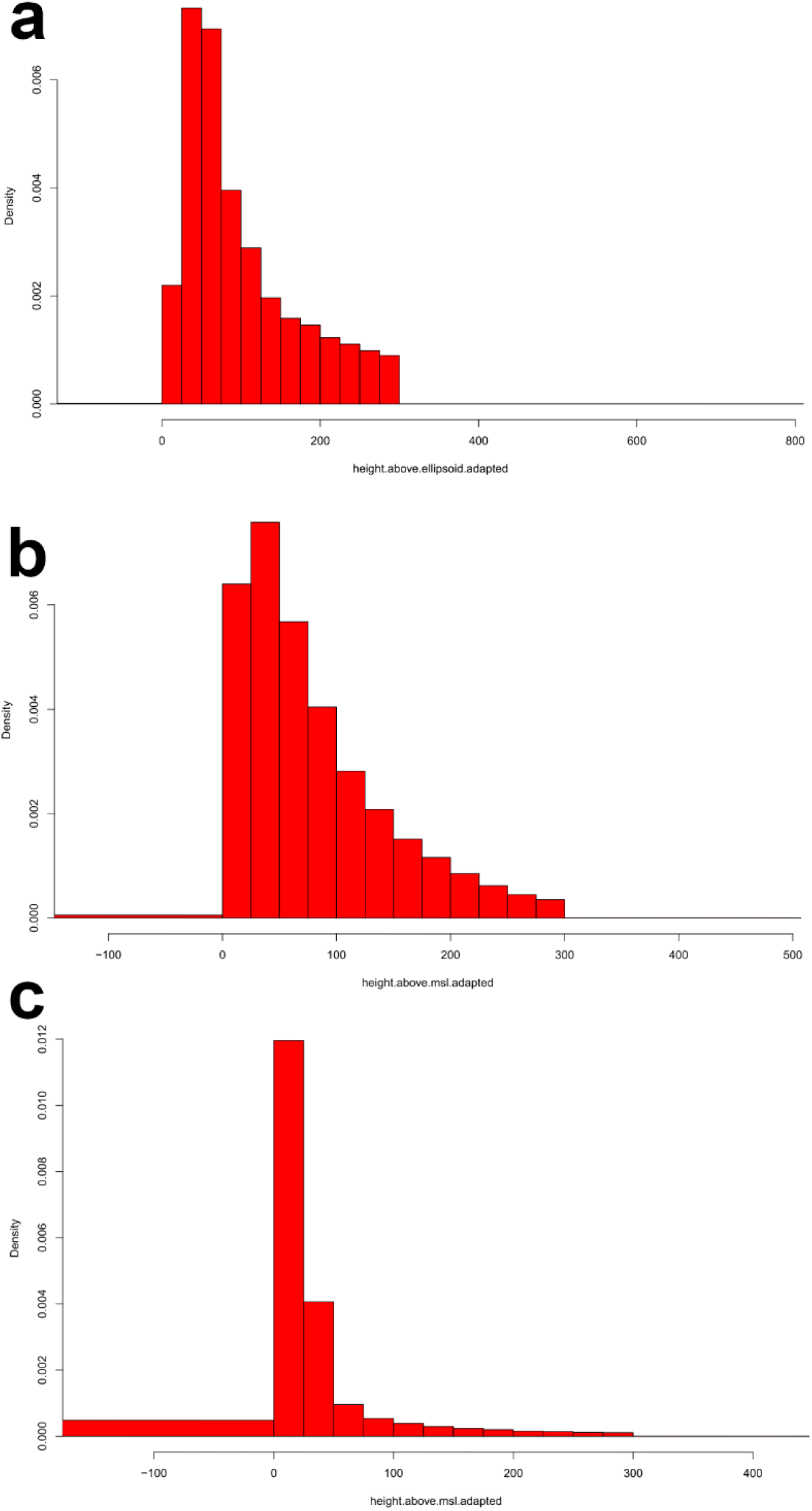
Histograms of average duration probability densities that tracked birds flew in windows of height above the ground between 0 and 300 m while nesting. (a) white stork, (b) red kite and (c) marsh harrier. Note that values below 0 and above 300 m were combined into one interval, respectively.

The final parameter, area usage in flight at different distances to the nest, also shows a skew towards the nest, especially the white stork showing a high peak below 100 m (Figure 2a). Still, the white storks use most distant areas of the three species, on average 3.1 km and up to 10 km from the nest (Table 2). Their 50 % usage threshold lies at 1400 m. Red kites show intermediate values with average use of 1.7 km and 50 % usage threshold at 1100 m. Marsh harries stay very close to their nest (average 1.1 km and 50 % usage threshold at 700 m) and show strong declines in use intensity with increasing distance to the nest (Figure 2c).

**Figure 2.**
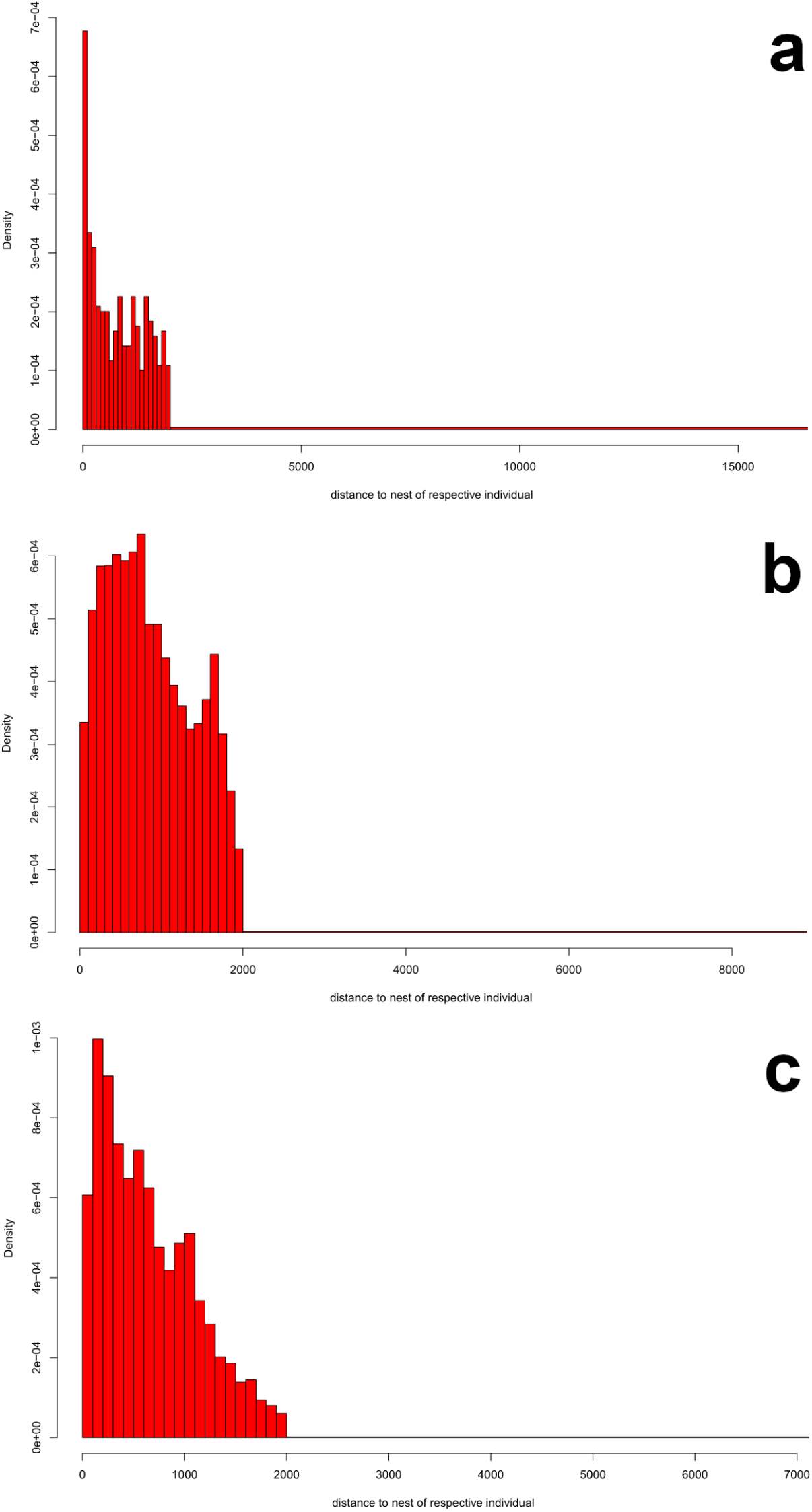
Histograms of average duration probability densities that tracked birds flew in different distance windows to the nest, between 0 and 2000 m. (a) white stork, (b) red kite and (c) marsh harrier. Note that values above 2000 m were combined into one interval.

## Discussion

We have developed a set of two workflows to estimate simple flight behaviour parameters during nesting times of large birds: flight speed, daily flight duration, flight height and distance to the nest when in flight. Including automated nest detection, these parameters are estimated from GPS tracks of nesting birds by the workflows in the interactive and easy-to-use, open analysis platform MoveApps with minimal required settings (Kölzsch et al. 2022). Because MoveApps is open source, our workflows are provided there as public workflows for use by any registered user, and they are available to anyone that needs to extract such values from tracking data. Furthermore, a link to Movebank allows the easy integration of other own or open tracking data sets (Kays et al. 2022). The parameters that can be extracted by our workflows provide insight into the behaviour and ecology of the respective birds and provide input parameters for collision risk models that are in use during wind turbine planning. Many individual and daily values are provided in the different App outcomes (see Appendices 2-4), so that results can be evaluated in much more detail than presented here. Our workflows have been developed in an unbiased manner from automatically collected location data, thus supporting the development of climate friendly energy generation as well as bird conservation.

While applying the developed workflows to three example species we became aware of several issues. First, our results reveal that nesting times of all three species are spread rather widely throughout the defined breeding season interval. Even if individual nesting durations are only some weeks long, the overall breeding season encompasses 5 months or longer. This is different from synchronously breeding, colonial birds (Emlen and Demong 1975), but known already for e.g. red kites (Mammen et al. 2013). The asynchrony might be either (1) adaptive or (2) timing of breeding events differs spatially by local conditions or between years due to climate differences (Marrot et al. 2018). This indicates that wind turbines next to nesting sites affect the birds during a long and unpredictable time interval each year.

Second, the detection of flight behaviour by tracking data can be approached in different ways – based on location height or speed. We have chosen for instantaneous GPS ground speed, as this measure shows the instantaneous velocity a bird moves with when its location is determined (Safi et al. 2013). This allows to also detect slow flights, as in soaring flight (Murgatroyd et al. 2021), which are important to consider around elevated nesting sites. The instantaneous property of this GPS ground speed makes it rather unaffected by low data resolution or gaps and flight height estimates have low variance within species. Flight duration, however is affected by data gaps. Therefore, we have calculated duration of flight based on the daily tracking duration. Any duration of gaps was not included and cannot be considered for the parameters. For true flight durations (per 24 h) we recommend additional GPS tracking studies with regular collection of locations (at least one location per 15-20 min), throughout the whole days and nights. Only then can unexpected flight behaviour during e.g. night be excluded or considered. In general, flight duration per day is rather low, especially for species that stay on the nest much and/or feed on the ground (white stork).

Third, flight heights as well as distances to the nest, which are the parameters that wind turbine placement can be adapted to mostly, show large variances between individuals of the same species. Both are likely mainly due to local conditions and habitat preferences, indicating that general parameters for a species must be set into local context before planning wind turbine construction sites. This can be done by including habitat properties and species preferences into the collision risk models and decision making processes (Krone and Treu 2018; Tikkanen et al. 2018). Furthermore, avoidance behaviour should be included, as nesting birds can adapt their flight behaviour around wind turbines (Watson et al. 2018), possibly leading to irregular habitat use.

Flight height as measured by GPS is generally not very accurate (Cleasby et al. 2015). With large data sets and averaging, this shortcoming can be accounted for, as is done in our examples. Further improvement could be gained by even higher data resolution (Schaub et al. 2020). Our average range of 50% flight altitude of red kites (50-75 m) is almost the same as previously estimated (50-60 m) for a data set in central Germany (Pfeiffer and Meyburg 2022). Furthermore, the issue of ellipsoid vs. mean sea level height at estimating height above ground has to be considered. For local applications as ours, this can be done manually, but more automated ways have to and will be developed. Meanwhile, conservative error bars should be drawn around estimated flight.

Nest distance seems more trustworthy with respect to location accuracy. Our 50% in-flight usage threshold for red kites (1100 m) coincides very closely with the previously calculated value of 1000 m from other tracking data (Mammen et al. 2013). For red kite and marsh harrier these values coincide very closely with the central evaluation area radii given in the Germany Federal Nature Conservation Act (Federal Law Gazette 2009): red kite: 1100 m vs. 1200 m, marsh harrier 700 m vs. 500 m. Only the white storks in our data set had a somewhat further usage area than required: 1400 m vs. 1000 m. When looking at individual’s GPS tracks on the map, irregular use of the landscape around nests become apparent. Birds have preferred sites to forage and often use dense corridors to reach those (Krone and Treu 2018). Thus, an improvement to our results by potentially including habitat selection and suitability (Tikkanen et al. 2018) is advisable. More recent collision risk models consider the circumstance of non-uniform area usage around the nest and include habitat properties surrounding the nesting and wind turbine site (BDEW Bundesverband der Energie-und Wasserwirtschaft e.V. 2021).

By the implementation of the here presented workflows in an open-source analysis platform like MoveApps, we not only alleviate the use of them for other data sets, but encourage the improvement and expansion of the code of its components that is openly available on GitHub. All code is written in R, which is an adaptive and continuously changing software. We strongly encourage the improvement of our Apps (see Appendix 1) as well as the modification of the workflows. In the above paragraphs, several avenues for improvement are discussed.

In conclusion, our workflows present a first step for a tracking-based analysis for wind turbine planning, providing insight into movement behaviour during nesting as well as simple, basic parameters for collision risk models. Evaluations of suitability have been performed here for three species, but more must be included. We have shown the high additional value of available tracking data sets that open up more possibilities to gain insight of animal behaviour in relation to wind turbines, not only during breeding, but potentially in winter and during possible migration movements (Bairlein 2016). MoveApps provides a suitable and available platform for additional development and improvements. We here provide a useful tool for initial indications of collision risk by wind turbines during nesting, alleviating one important threat to birds and other flying animals. Our tool is freely available and we look forward to it being widely used, adapted and expanded by the community for providing the best parameters and statistics for safe wind turbine planning.

## Acknowledgements

We are grateful for discussion and comments with Kolja Wolanska and Bettina Wilkening and funding that supported the development of the workflows by ENERTRAG SE and wpd onshore GmbH & Co. KG.

We recognise funding from the “Landesanstalt für Umwelt Baden-Württemberg” for tracking the red kites. Data of marsh harriers has been provided via Tonio Schaub and Peter Desmet by the Belgian Research Institute for Nature and Forest (INBO) and data of white storks by the Max Planck Institute of Animal Behavior. White stork nest detection was done with help by Andrea Flack. We are grateful to Wolfgang Fiedler, Kamran Safi and Tonio Schaub for discussions about the workflows and comments on a previous version of this manuscript.

## Data and Code Availability

The white stork data set as well as the marsh harrier data are available as public studies in Movebank (see Methods). The red kite data set is also stored in Movebank, and access can be granted upon reasonable request to the corresponding author.

Both developed workflows are public workflows in MoveApps (moveapps.org) and can be used there. They are also published in the Movebank Data Repository: Nest Location and Nesting Duration Workflow https://doi.org/10.5441/001/1.4473qv78 (Kölzsch and Flack 2022) and Nest Flight Statistics Workflow https://doi.org/10.5441/001/1.v45k1d06 (Kölzsch and Gal 2022).

## Appendix 1 Workflow Vignette

### 1 Nest Location and Nesting Duration Workflow

In the Nest Detection workflow (Figure A1), first the data from Movebank are loaded, in this case preferably with a resolution not higher than 1 location per hour, because many distance calculations are required. Higher resolutions might lead to a crash of the Nest Cluster Detection App. For this workflow, only full breeding seasons are of interest, therefore only tracks of animals with at least data of one full breeding season should be loaded. In the next step, each track is split into single years/seasons. All further calculations are done on those single year tracks.

The Nest Cluster Detection App is the centre of this workflow. It needs a number of input parameters (Settings) regarding the nesting duration, percentage of nest attendance etc., that should ideally be evaluated by species experts. For details about the algorithm see also the documentation of the underlying R package nestR (Picardi et al. 2020). The App output includes all locations of the data set’s tracks during nesting attempts and can be visualised with e.g. the Interactive Map (leaflet) App.

**Figure A1.**
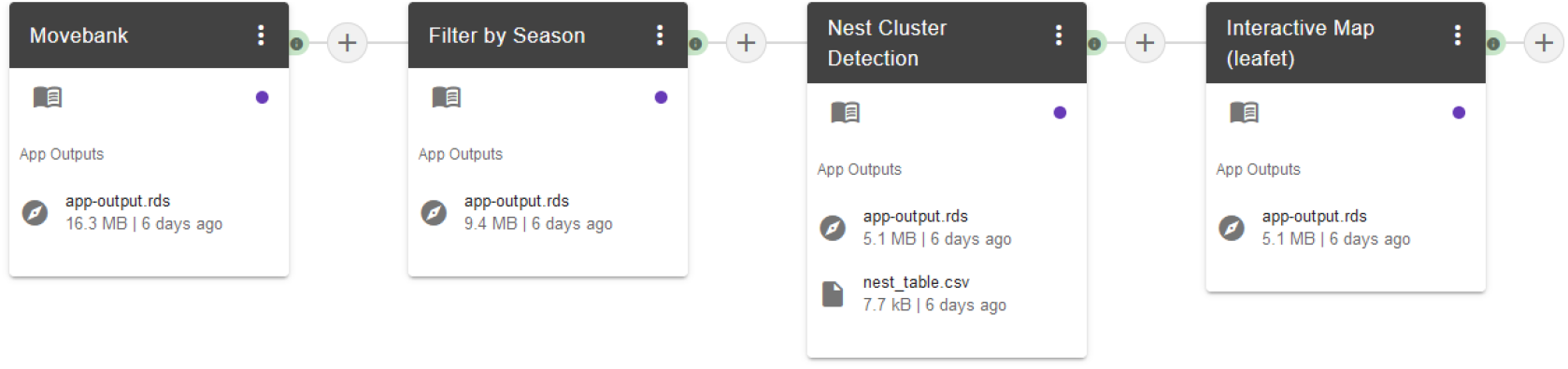
Nest Location and Nesting Duration Workflow as shown in MoveApps.

The nest_table.csv that is created by the Nest Cluster Detection App is the main output of this App. For each track, it contains the nest location, nest visit start and end times and properties of nest attendance and revisitation. This information can be used independently, but it is also important input for the second workflow below and needs to be evaluated once before, manually. Please go through it and check if the nesting attempts are sensible. Else, either adapt the App Settings (and rerun) or manually select and adapt sensible rows. In the latter case, make sure that the format of the columns is not changed. For use in the second workflow, please name this file by a convenient name and add it to your Cloud Storage folder that is linked with MoveApps.

If nest use is known from observations, it is also possible to prepare a similar file for use in the below, second workflow. Then, it is important to keep the following columns in the file with the related data formats: individual.local.identifier, timestamp, location.long, location.lat, individual.taxon.canonical.name, sensor.type.

### 2 Nest Flight Statistics Workflow

Based on the detected (or observed) nesting sites and time intervals of nest use, the second workflow (Figure A2) calculates parameters of flight and nest use behaviour that can be used as input for wind turbine planning models. The nest_table.csv of the first workflow must be added to the data analysis at two points of the workflow: (i) before the Filter by Individual Time Intervals App to only analyse the individual nesting intervals for each track and (ii) before the Radial Area Use Around Location App to provide nest locations for distance calculations and plots.

The workflow again starts with downloading tracks from Movebank. For the following calculations, high resolution data is necessary, at least 1 location per 15 minutes. Therefore, filtering like in the first workflow should not be done, but rather to at most 1 location per 1 minute if there are bursts in the data (that are not of interest for the analyses here). For combining this workflow with the nest detection results of the first workflow, it is important that in the download step exactly the same individuals and time ranges are selected as there. Only the data resolution should differ. Thus, also the Filter by Season App has to come next with the same settings as in the first workflow.

Next, the Upload File from Cloud Storage App is uploading the nest_table.csv from the to your account linked Cloud Storage folder (either Dropbox or Google Drive) and appending it to the data for being used in the Filter by Individual Time Intervals App. The output of the latter App is again only the data without the nest details table information, but filtered by the individual-specific nesting intervals.

**Figure A2.**
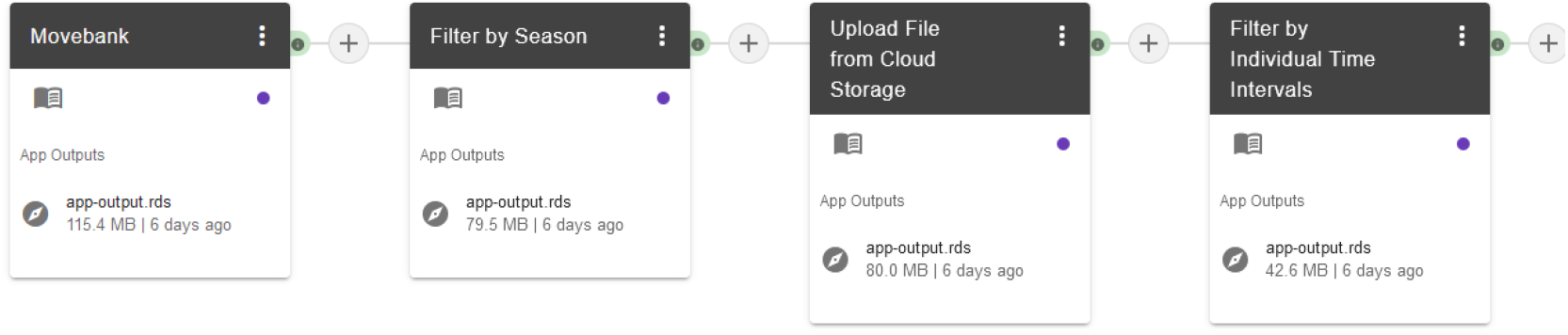
First part of the Nest Flight Statistics Workflow as shown in MoveApps. It entails data upload and cleaning.

#### 2.1 Flight speed

The Extract Two Movement Speeds App plots histograms of GPS ground speed and fits a bimodal distribution to it (Figure A3). The attribute GPS ground speed, called “ground.speed” in the Workflow, must be in your data! It is the GPS collected instantaneous speed of the tracked animal. Inter-location speed estimations should not be used to approximate flight speed in this App. The output parameters in the groundspeed_modes.csv file are mode1 (average speed of non-flight), antimode (turning point between the two behaviours) and mode2 (average speed of flight). For some of the following Apps the average antimode can be used as threshold ground speed (manually add to Settings) to filter for flight behaviour. Histograms of ground speed for each track are provided as pdf output.

**Figure A3.**
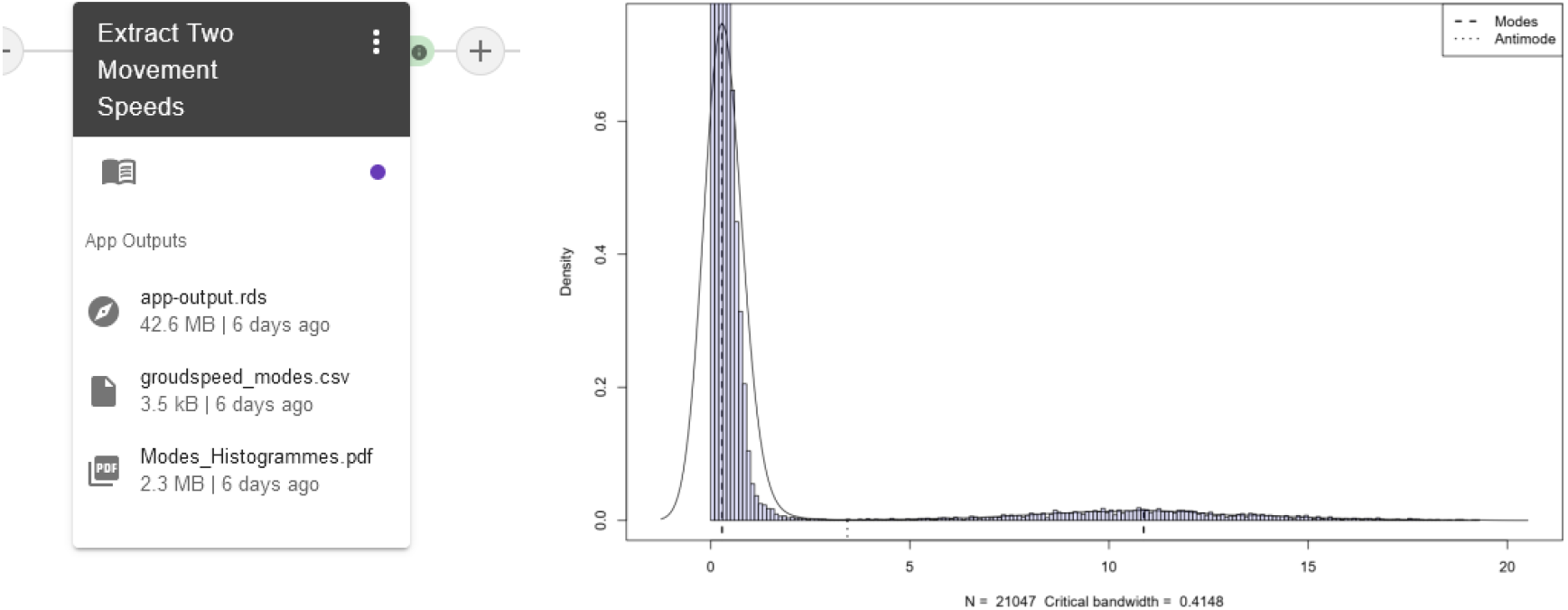
Extract Two Movement Speeds App as part of the second MoveApps Workflow. On the right, see a histogramme of GPS ground speed of one animal with the fitted bimodal distribution.

#### 3.2. Daily flight duration

The duration of birds flying every day during the nesting season indicates how often they might come close to wind turbines in the region. Therefore, the following Apps (Figure A4) filter the data and calculate the proportion of locations in flight per day for each individual and day and average them. As the proportion of locations might be biased by unregular data collection, we additionally consider location duration, i.e. each location is attributed with the duration until the next location, assuming that the same behaviour (flight/no flight) is retained for that duration. This duration is appended to each location by the Time Lag Between Locations App. Consider adapting the time unit to hours for further analyses.

Furthermore, if the tags were systematically set to not collect data at night (i.e. because it is expected that the birds stay at their nest then), the calculation using duration can become biased if the last locations of the day are flight, because they are attributed with a very long duration. Also, other gaps can be present in the data, e.g. due to lack of energy or no visibility of GPS satellites. Using the Adapt/Filter Unrealistic Values App, locations with such occurring long durations can be adapted or taken out from the analysis. We advise to provide a realistic threshold value and set the time lag of the respective locations to the median of all others.

Finally, the Daily Proportions App determines the proportion of flight behaviour per daily tracking duration (note that this is often less than the 24h). By multiplication of flight proportion and daily tracking duration, one can obtain daily flight duration in the required time interval. Flight is here defined as moving with a ground speed above a defined threshold. This threshold should be set close to the average antimode obtained by the Extract Two Movement Speeds App.

**Figure A4.**
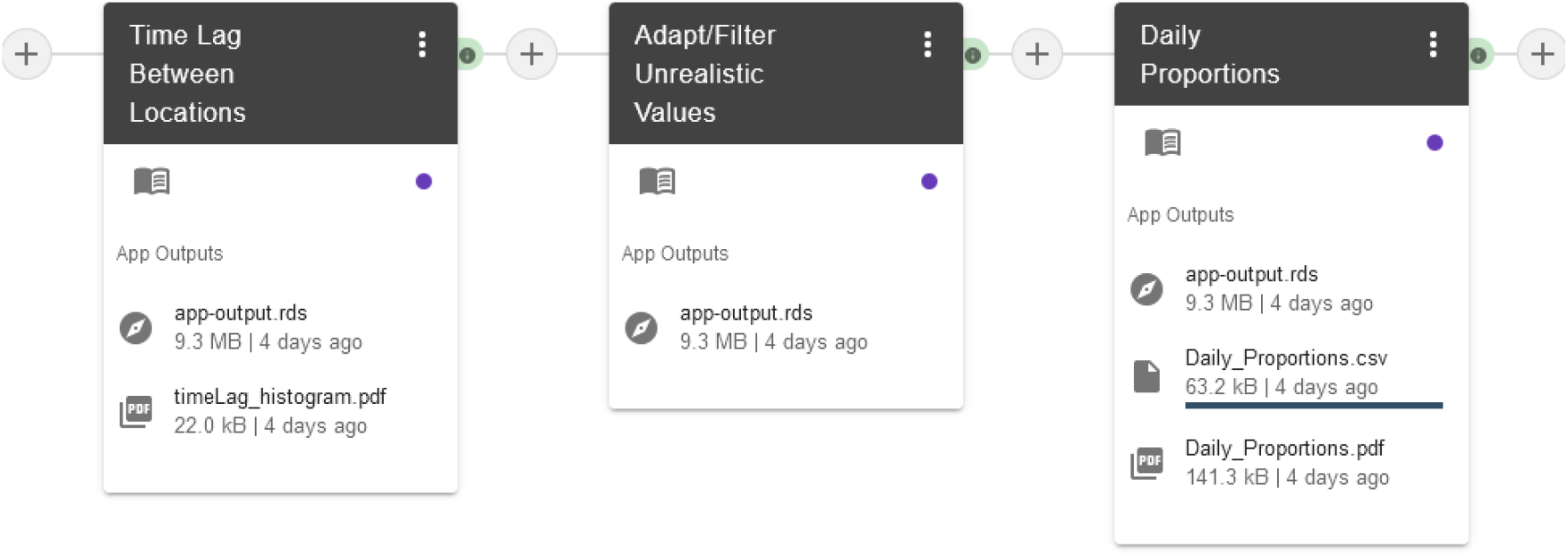
Daily Proportions App for determination of daily flight durations and preparations for it as part of the second MoveApps Workflow.

#### 3.3 Flight height

Depending on the wind turbine’s height, flight height is important to estimate how often the birds fly in the dangerous heights of the turning blades. For the following analyses (Figure A5), only locations in flight are of interest, so that the Filter by Speed App is used to keep only locations with ground speed above the estimated flight speed antimode.

GPS tags collect, depending on the model and company, height above mean sea level or height above the earth ellipsoid. To convert this into height above ground, one has to adapt the height variable with ground heights. These are provided globally by Digital Elevation Models. Using the R package elevatr, the Add Elevation and Height above Ground App is appending each location with this ground elevation and is adapting available height by subtraction.

If the tags collect height above mean sea level (often called height.above.msl), the adapted value is accurate to a few metres. However, if height.above.ellipsoid is calculated, the inaccuracy is larger, in the range of 50-100 m. If higher accuracies are necessary, a transformation is advised. As the transformation between height.above.msl and height.above.ellipsoid is complex and differs strongly by region on the globe, we have so for not included this transformation into the workflow. It is planned for a future version under the use of the Earth Gravitational Model (https://earth-info.nga.mil/index.php?dir=wgs84&action=wgs84#tab_egm2008). For analyses in Germany it can be said that in general, mean sea level is 30-40 m above the ellipsoid, so that we added 30 m to adapted ellipsoid height for the true ground height. In addition to average flight heights, aggregated values of proportional use of flight height are provided, always cumulative between the selected height and the ground.

**Figure A5.**
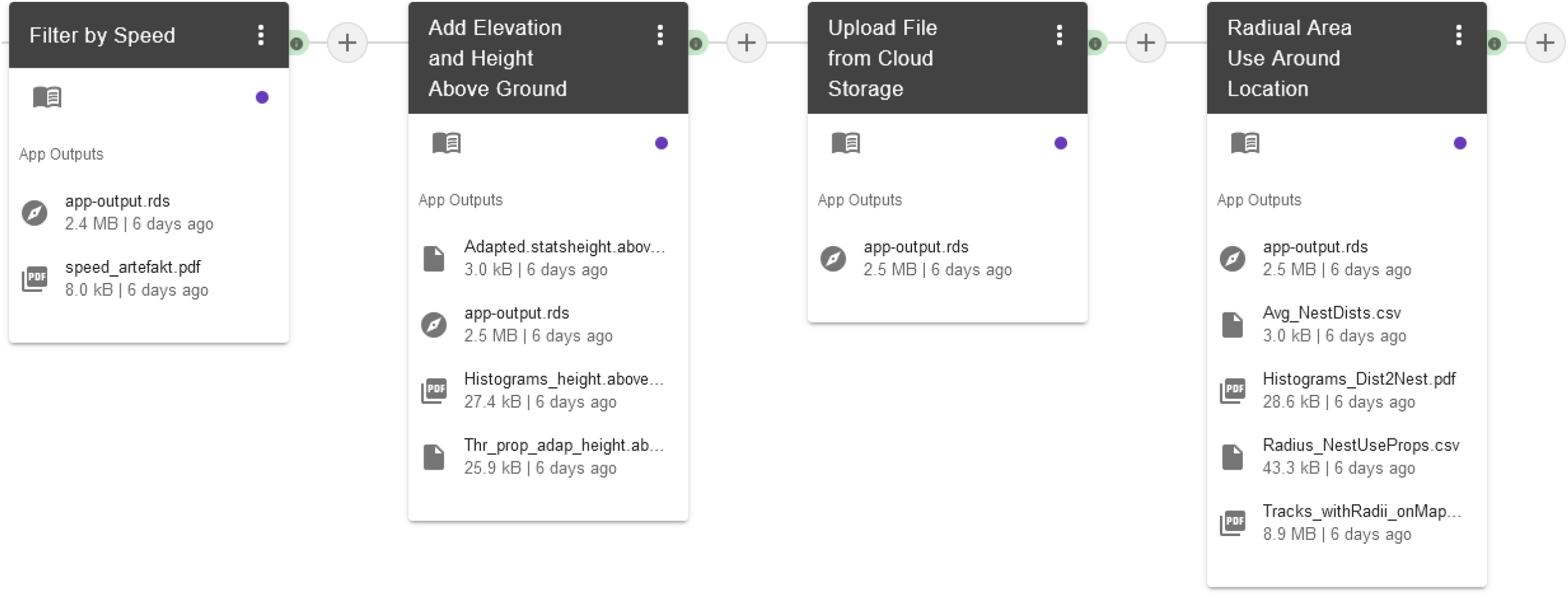
Final part of the second MoveApps workflow, including flight height and distance to the nest.

#### 3.4 Distance to nest

To evaluate how far from a nest wind turbines can be erected, proportional use of the area around the nest site must be evaluated. This parameter has been calculated in some species-specific studies and is related to central evaluation area radii given in the German Federal Nature Conservation Act (Federal Law Gazette 2009), so it can be compared with outcomes of this part of the workflow. The Radial Area Use Around Location App calculates cumulatively to the nest the proportion of area use until a set of circular distances around the nest. For visual support a histogram as well as a map with the radii around the nest is provided, but it must be made clear that this plot does not show any results like use kernels. However, from the maps it becomes clear that usage of the area around the nest is not uniform, but likely dependent on forage availability and corridors to areas of good foraging.

## Appendix 2 Individual results for white storks

**Table A1.** Individual nesting times and durations for the selected white stork tracks per year as extracted from the Nest Cluster Detection App.

| burst | first_date | last_date | attempt_start | attempt_end | days_vis | visit_dur | attempt_dur |
| --- | --- | --- | --- | --- | --- | --- | --- |
| Angela...DER.AY470..eobs.4001.6_2021 | 2021-03-01 | 2021-08-31 | 2021-03-02 | 2021-07-29 | 181 | 183 | 149 |
| Angela...DER.AY470..eobs.4001.7_2022 | 2022-03-01 | 2022-07-31 | 2022-03-01 | 2022-07-28 | 146 | 152 | 149 |
| Europa...DER.A1A26..eobs.4004.4_2019 | 2019-03-20 | 2019-08-31 | 2019-03-20 | 2019-08-16 | 116 | 164 | 149 |
| Europa...DER.A1A26..eobs.4004.5_2020 | 2020-03-01 | 2020-07-19 | 2020-03-01 | 2020-07-19 | 134 | 140 | 140 |
| Europa...DER.A1A26..eobs.4004.6_2021 | 2021-03-04 | 2021-08-31 | 2021-03-31 | 2021-08-27 | 172 | 180 | 149 |
| Europa...DER.A1A26..eobs.4004.7_2022 | 2022-03-01 | 2022-08-31 | 2022-03-01 | 2022-07-28 | 184 | 183 | 149 |
| Louis...DER.AU050..eobs.3264.3_2018 | 2018-04-08 | 2018-08-31 | 2018-04-08 | 2018-08-31 | 146 | 145 | 145 |
| Louis...DER.AU050..eobs.3264.4_2019 | 2019-03-01 | 2019-08-23 | 2019-03-01 | 2019-07-28 | 176 | 175 | 149 |
| Louis...DER.AU050..eobs.3264.5_2020 | 2020-03-01 | 2020-08-12 | 2020-03-01 | 2020-07-28 | 165 | 164 | 149 |
| Louis...DER.AU050..eobs.3264.6_2021 | 2021-03-01 | 2021-08-31 | 2021-03-08 | 2021-08-04 | 184 | 183 | 149 |
| Louis...DER.AU050..eobs.3264.7_2022 | 2022-03-01 | 2022-08-23 | 2022-03-10 | 2022-08-06 | 134 | 175 | 149 |
| Mirabell.....DER.AN910..eobs.3907.3_2018 | 2018-04-05 | 2018-08-26 | 2018-04-05 | 2018-08-26 | 143 | 143 | 143 |
| Mirabell.....DER.AN910..eobs.3907.4_2019 | 2019-03-17 | 2019-08-30 | 2019-03-18 | 2019-08-14 | 167 | 166 | 149 |
| Mirabell.....DER.AN910..eobs.3907.5_2020 | 2020-03-16 | 2020-08-31 | 2020-03-17 | 2020-08-13 | 169 | 168 | 149 |
| Niclas...DER.AU053..eobs.3341.1_2016 | 2016-03-10 | 2016-08-28 | 2016-03-11 | 2016-08-07 | 153 | 171 | 149 |
| Sara.....DER.AU017..eobs.3941.2_2017 | 2017-03-05 | 2017-06-12 | 2017-03-05 | 2017-06-12 | 96 | 99 | 99 |
| Sara.....DER.AU017..eobs.3941.3_2018 | 2018-03-09 | 2018-08-05 | 2018-03-09 | 2018-08-05 | 149 | 149 | 149 |
| Sara.....DER.AU017..eobs.3941.4_2019 | 2019-03-11 | 2019-08-25 | 2019-03-12 | 2019-08-08 | 155 | 167 | 149 |
| Sierit....DER.AN858..eobs2561.1_2016 | 2016-04-17 | 2016-08-30 | 2016-04-20 | 2016-08-30 | 128 | 135 | 132 |
| Sierit....DER.AN858..eobs2561.3_2018 | 2018-03-17 | 2018-08-31 | 2018-03-20 | 2018-08-16 | 165 | 167 | 149 |
| Sierit....DER.AN858..eobs2561.4_2019 | 2019-03-21 | 2019-08-31 | 2019-05-05 | 2019-08-31 | 125 | 163 | 118 |
| Sierit....DER.AN858..eobs2561.5_2020 | 2020-03-07 | 2020-08-31 | 2020-03-07 | 2020-08-03 | 178 | 177 | 149 |
| Sierit....DER.AN858..eobs2561.6_2021 | 2021-03-04 | 2021-08-31 | 2021-03-05 | 2021-08-01 | 181 | 180 | 149 |
| Zozu.....DER.AL581..eobs2541.1_2016 | 2016-03-21 | 2016-08-30 | 2016-03-22 | 2016-08-18 | 162 | 162 | 149 |
| **Zozu.....DER.AL581..eobs2541.2_2017** | 2017-03-04 | 2017-08-28 | 2017-03-05 | 2017-08-01 | 165 | 177 | 149 |
| **Zozu.....DER.AL581..eobs2541.3_2018** | 2018-03-05 | 2018-08-21 | 2018-03-05 | 2018-08-01 | 161 | 169 | 149 |
| **Zozu.....DER.AL581..eobs2541.4_2019** | 2019-03-09 | 2019-08-21 | 2019-03-09 | 2019-08-05 | 165 | 165 | 149 |
| **Zozu.....DER.AL581..eobs2541.5_2020** | 2020-03-05 | 2020-08-20 | 2020-03-05 | 2020-08-01 | 154 | 168 | 149 |

**Table A2.** Individual flight speed estimates for the selected white stork tracks per year as extracted from the Extract Two Movement Speeds App.

| **trackId** | **mode1** | **antimode** | **mode2** | **mean.above.antimode** | **sd.above.antimode** |
| --- | --- | --- | --- | --- | --- |
| **Mirabell.....DER.AN910..eobs.3907.3** | 0.243965124 | 3.638685964 | 11.20703765 | 10.4194571 | 2.4040619 |
| **Mirabell.....DER.AN910..eobs.3907.4** | 0.206598897 | 3.735720966 | 10.06768554 | 10.2213277 | 2.58274334 |
| **Mirabell.....DER.AN910..eobs.3907.5** | 0.228510841 | 4.689618198 | 10.90902698 | 10.461164 | 2.27034618 |
| **Angela...DER.AY470..eobs.4001.6** | 0.310842225 | 3.715733428 | 11.61837189 | 10.7781168 | 2.87455014 |
| **Angela...DER.AY470..eobs.4001.7** | 0.28766587 | 3.432550922 | 10.86977909 | 10.8559597 | 2.89513522 |
| **Europa...DER.A1A26..eobs.4004.4** | 0.222912616 | 4.00517304 | 10.96835584 | 11.2660176 | 2.93608145 |
| **Europa...DER.A1A26..eobs.4004.5** | 0.24186948 | 4.431192721 | 12.05509226 | 11.1985991 | 2.97741063 |
| **Europa...DER.A1A26..eobs.4004.6** | 0.217180583 | 4.504138157 | 11.29544813 | 11.3397766 | 2.84610515 |
| **Europa...DER.A1A26..eobs.4004.7** | 0.252317434 | 4.019572804 | 11.10854743 | 11.2997285 | 2.96071973 |
| **Louis...DER.AU050..eobs.3264.3** | 0.201612822 | 4.388988558 | 10.05601684 | 10.477933 | 2.36920222 |
| **Louis...DER.AU050..eobs.3264.4** | 0.218390101 | 4.183016008 | 10.43240567 | 10.501327 | 2.54640073 |
| **Louis...DER.AU050..eobs.3264.5** | 0.223607892 | 3.91016217 | 10.46496694 | 10.241602 | 2.58743335 |
| **Louis...DER.AU050..eobs.3264.6** | 0.200182076 | 3.917676676 | 10.87187744 | 10.2683688 | 2.36669446 |
| **Louis...DER.AU050..eobs.3264.7** | NA | NA | NA | NA | NA |
| **Niclas...DER.AU053..eobs.3341.1** | 0.264702107 | 4.879529535 | 10.34702769 | 10.8958654 | 2.35435555 |
| **Sara.....DER.AU017..eobs.3941.2** | 0.199297317 | 3.388701331 | 10.10558932 | 11.0837821 | 3.15048704 |
| **Sara.....DER.AU017..eobs.3941.3** | 0.179665785 | 3.768918872 | 10.52245214 | 10.9772685 | 3.01973427 |
| **Sara.....DER.AU017..eobs.3941.4** | 0.220358703 | 3.436448433 | 10.74356325 | 10.6528222 | 3.03089906 |
| **Sierit....DER.AN858..eobs2561.1** | 0.283838825 | 5.725754609 | 11.08440895 | 10.9485377 | 2.41752031 |
| **Sierit....DER.AN858..eobs2561.3** | 0.359728359 | 5.382840046 | 11.00743081 | 11.1035348 | 2.90718204 |
| **Sierit....DER.AN858..eobs2561.4** | 0.297934231 | 5.943178207 | 11.65163747 | 11.2577465 | 2.24893446 |
| **Sierit....DER.AN858..eobs2561.5** | 0.321308223 | 5.957681408 | 11.17518627 | 10.8525676 | 2.45060382 |
| Sierit....DER.AN858..eobs2561.6 | 0.278558565 | 5.866233539 | 11.72416441 | 10.7531456 | 2.2757734 |
| Zozu.....DER.AL581..eobs2541.1 | 0.279937082 | 6.422220221 | 11.16558913 | 11.3311436 | 2.30427716 |
| Zozu.....DER.AL581..eobs2541.2 | 0.323213973 | 5.33459777 | 11.02356893 | 11.2332771 | 2.80200789 |
| Zozu.....DER.AL581..eobs2541.3 | 0.307218405 | 5.169435081 | 10.87006814 | 11.0887335 | 2.78030428 |
| Zozu.....DER.AL581..eobs2541.4 | 0.306940625 | 5.800542493 | 11.36277401 | 11.5940906 | 2.88398728 |
| Zozu.....DER.AL581..eobs2541.5 | 0.32512918 | 4.70852714 | 10.34576904 | 11.0680639 | 3.07049575 |
| mean | 0.25938842 | 4.605808826 | 10.92792005 | 10.8951836 | 2.67827581 |
| sd | 0.048807602 | 0.921321238 | 0.519622688 | 0.38155218 | 0.29532111 |

**Table A3.** Individual daily flight duration estimates for the selected white stork tracks per year as extracted from the Daily Proportions App.

| trackId | date | n.pts | perc_pts | perc_dur | track_dur |
| --- | --- | --- | --- | --- | --- |
| Mirabell.....DER.AN910..eobs.3907.3 | avg | 147 | 0.034613636 | 0.035010429 | 17.92702991 |
| Mirabell.....DER.AN910..eobs.3907.4 | avg | 135 | 0.022507711 | 0.022627458 | 17.99965697 |
| Mirabell.....DER.AN910..eobs.3907.5 | avg | 149 | 0.024549991 | 0.02453441 | 17.99874161 |
| Angela...DER.AY470..eobs.4001.6 | avg | 140 | 0.048267126 | 0.046149531 | 16.90626984 |
| Angela...DER.AY470..eobs.4001.7 | avg | 149 | 0.082846366 | 0.082535108 | 17.72057819 |
| Europa...DER.A1A26..eobs.4004.4 | avg | 134 | 0.067814252 | 0.068162486 | 17.99958613 |
| Europa...DER.A1A26..eobs.4004.5 | avg | 145 | 0.042945425 | 0.043681464 | 17.63797619 |
| Europa...DER.A1A26..eobs.4004.6 | avg | 149 | 0.050865154 | 0.050402874 | 17.96405481 |
| Europa...DER.A1A26..eobs.4004.7 | avg | 149 | 0.076850119 | 0.078149345 | 15.15353027 |
| Louis...DER.AU050..eobs.3264.3 | avg | 149 | 0.058292225 | 0.059534114 | 17.99885249 |
| Louis...DER.AU050..eobs.3264.4 | avg | 122 | 0.04209136 | 0.04174318 | 17.94832774 |
| Louis...DER.AU050..eobs.3264.5 | avg | 143 | 0.040552189 | 0.04058038 | 17.96693326 |
| Louis...DER.AU050..eobs.3264.6 | avg | 149 | 0.032829292 | 0.033216738 | 17.96420022 |
| Louis...DER.AU050..eobs.3264.7 | avg | 149 | 0 | 0 | 0.084016393 |
| Niclas...DER.AU053..eobs.3341.1 | avg | 149 | 0.044728647 | 0.044842605 | 17.9825522 |
| Sara.....DER.AU017..eobs.3941.2 | avg | 96 | 0.085853039 | 0.076901902 | 15.33159722 |
| Sara.....DER.AU017..eobs.3941.3 | avg | 149 | 0.069115088 | 0.064525518 | 14.70660887 |
| Sara.....DER.AU017..eobs.3941.4 | avg | 149 | 0.058360241 | 0.04951458 | 13.20362975 |
| Sierit....DER.AN858..eobs2561.1 | avg | 132 | 0.032401846 | 0.033247071 | 16.61925076 |
| Sierit....DER.AN858..eobs2561.3 | avg | 149 | 0.040018149 | 0.04069823 | 17.99672066 |
| Sierit....DER.AN858..eobs2561.4 | avg | 118 | 0.016444661 | 0.017028573 | 18.00009417 |
| Sierit....DER.AN858..eobs2561.5 | avg | 149 | 0.035515921 | 0.034864142 | 17.98942394 |
| Sierit....DER.AN858..eobs2561.6 | avg | 149 | 0.027111667 | 0.02773887 | 17.96693514 |
| Zozu.....DER.AL581..eobs2541.1 | avg | 149 | 0.028601219 | 0.029215294 | 18.00031695 |
| Zozu.....DER.AL581..eobs2541.2 | avg | 149 | 0.031520278 | 0.031971011 | 11.8656115 |
| Zozu.....DER.AL581..eobs2541.3 | avg | 149 | 0.035953903 | 0.035961834 | 17.98268834 |
| Zozu.....DER.AL581..eobs2541.4 | avg | 149 | 0.045859473 | 0.047076989 | 17.97636838 |
| Zozu.....DER.AL581..eobs2541.5 | avg | 149 | 0.038719633 | 0.040035502 | 17.97889821 |
| mean | avg | 3994 | 0.043120115 | 0.042665226 | 16.64116696 |
| sd | avg | 3994 | 0.047148349 | 0.045304238 | 4.397497162 |

**Table A4.** Individual flight height estimates (average by number of points or duration) for the selected white stork tracks per year as extracted from the Add Elevation and Height Above Ground App.

| trackId | n.pts | mean.pts.height.adapted | sd.pts.height.adapted | mean.dur.height.adapted | sd.dur.height.adapted |
| --- | --- | --- | --- | --- | --- |
| Mirabell.....DER.AN910..eobs.3907.3 | 1067 | 110.4523899 | 150.3083844 | 110.36444 | 148.7965577 |
| Mirabell.....DER.AN910..eobs.3907.4 | 723 | 97.09488243 | 164.6821446 | 94.52381641 | 144.5986675 |
| Mirabell.....DER.AN910..eobs.3907.5 | 789 | 82.01356147 | 107.8847688 | 81.45900105 | 107.4481481 |
| Angela...DER.AY470..eobs.4001.6 | 1152 | 178.7963542 | 225.5804938 | 164.5712008 | 242.2438584 |
| Angela...DER.AY470..eobs.4001.7 | 2071 | 183.1622887 | 231.4910376 | 178.8441839 | 277.5082222 |
| Europa...DER.A1A26..eobs.4004.4 | 2183 | 181.6283097 | 178.6623953 | 181.0158697 | 179.1870019 |
| Europa...DER.A1A26..eobs.4004.5 | 1106 | 209.3339964 | 237.5571353 | 197.629241 | 224.082904 |
| Europa...DER.A1A26..eobs.4004.6 | 1504 | 189.1797207 | 352.825983 | 188.1095271 | 364.427967 |
| Europa...DER.A1A26..eobs.4004.7 | 2040 | 212.1713235 | 163.0415553 | 207.4181556 | 162.6356643 |
| Louis...DER.AU050..eobs.3264.3 | 1817 | 116.4577325 | 186.4350876 | 116.0807682 | 188.8884908 |
| Louis...DER.AU050..eobs.3264.4 | 1288 | 104.050854 | 351.5917407 | 101.2844645 | 347.6130621 |
| Louis...DER.AU050..eobs.3264.5 | 1219 | 87.94840033 | 193.5148485 | 85.77280367 | 186.2899515 |
| Louis...DER.AU050..eobs.3264.6 | 997 | 76.90882648 | 329.1519457 | 75.68891097 | 331.7840535 |
| Niclas...DER.AU053..eobs.3341.1 | 1392 | 142.6107759 | 229.8593529 | 143.6632307 | 226.8460433 |
| Sara.....DER.AU017..eobs.3941.2 | 933 | 212.2378349 | 195.0438339 | 195.2950673 | 185.3381213 |
| Sara.....DER.AU017..eobs.3941.3 | 1964 | 234.4559063 | 219.1148545 | 235.5179138 | 220.1595248 |
| Sara.....DER.AU017..eobs.3941.4 | 1071 | 197.8095238 | 164.5082156 | 195.4166189 | 163.2541706 |
| Sierit....DER.AN858..eobs2561.1 | 905 | 130.1055249 | 571.3632125 | 131.434354 | 600.7535109 |
| Sierit....DER.AN858..eobs2561.3 | 1241 | 225.6146656 | 1151.259006 | 226.9345134 | 1155.337684 |
| Sierit....DER.AN858..eobs2561.4 | 405 | 152.957037 | 1159.909428 | 132.4872506 | 1022.968448 |
| Sierit....DER.AN858..eobs2561.5 | 1057 | 131.1505203 | 258.0930348 | 127.4429697 | 252.2558041 |
| Sierit....DER.AN858..eobs2561.6 | 810 | 124.7225926 | 1182.982673 | 115.9655161 | 1089.956663 |
| Zozu.....DER.AL581..eobs2541.1 | 904 | 131.7120575 | 284.088899 | 127.2145024 | 274.0852711 |
| Zozu.....DER.AL581..eobs2541.2 | 987 | 184.9375887 | 445.9882496 | 182.0499717 | 473.7416599 |
| Zozu.....DER.AL581..eobs2541.3 | 1099 | 240.6118289 | 1892.643993 | 228.8693139 | 1808.831763 |
| Zozu.....DER.AL581..eobs2541.4 | 1393 | 200.7089734 | 839.3081912 | 195.1683424 | 836.8596579 |
| Zozu.....DER.AL581..eobs2541.5 | 1159 | 172.7089733 | 408.9575545 | 161.530518 | 388.4505367 |
| all | 33276 | 166.0934698 | 557.4652306 | 162.5549943 | 544.5013878 |

**Table A5.** Individual flight height aggregates for the selected white stork tracks per year as extracted from the Add Elevation and Height Above Ground App.

| trackId | height_threshold | n.loc | prop.locs | prop.dur |
| --- | --- | --- | --- | --- |
| mean | 25 | 27 | 0.088726674 | 0.089909801 |
| mean | 50 | 27 | 0.297721534 | 0.300012186 |
| mean | 75 | 27 | 0.477896895 | 0.486442822 |
| mean | 100 | 27 | 0.573197852 | 0.583265706 |
| mean | 125 | 27 | 0.638454095 | 0.648554787 |
| mean | 150 | 27 | 0.683610971 | 0.692920308 |
| mean | 175 | 27 | 0.720160368 | 0.728940432 |
| mean | 200 | 27 | 0.753525282 | 0.761886987 |
| mean | 225 | 27 | 0.781133301 | 0.789213109 |
| mean | 250 | 27 | 0.805847417 | 0.812700325 |
| mean | 275 | 27 | 0.828043962 | 0.834450242 |
| **mean** | 300 | 27 | 0.848170603 | 0.853862098 |
| **sd** | 25 | 27 | 0.075141504 | 0.078772031 |
| **sd** | 50 | 27 | 0.203825985 | 0.206376613 |
| **sd** | 75 | 27 | 0.193407836 | 0.191580302 |
| **sd** | 100 | 27 | 0.170096012 | 0.167507834 |
| **sd** | 125 | 27 | 0.141203959 | 0.137490179 |
| **sd** | 150 | 27 | 0.126267594 | 0.123202859 |
| **sd** | 175 | 27 | 0.114444841 | 0.110799155 |
| **sd** | 200 | 27 | 0.104705794 | 0.100886766 |
| **sd** | 225 | 27 | 0.095443073 | 0.091300842 |
| **sd** | 250 | 27 | 0.086182271 | 0.08263811 |
| **sd** | 275 | 27 | 0.076463083 | 0.072914346 |
| **sd** | 300 | 27 | 0.069411524 | 0.06607095 |

**Table A6.**
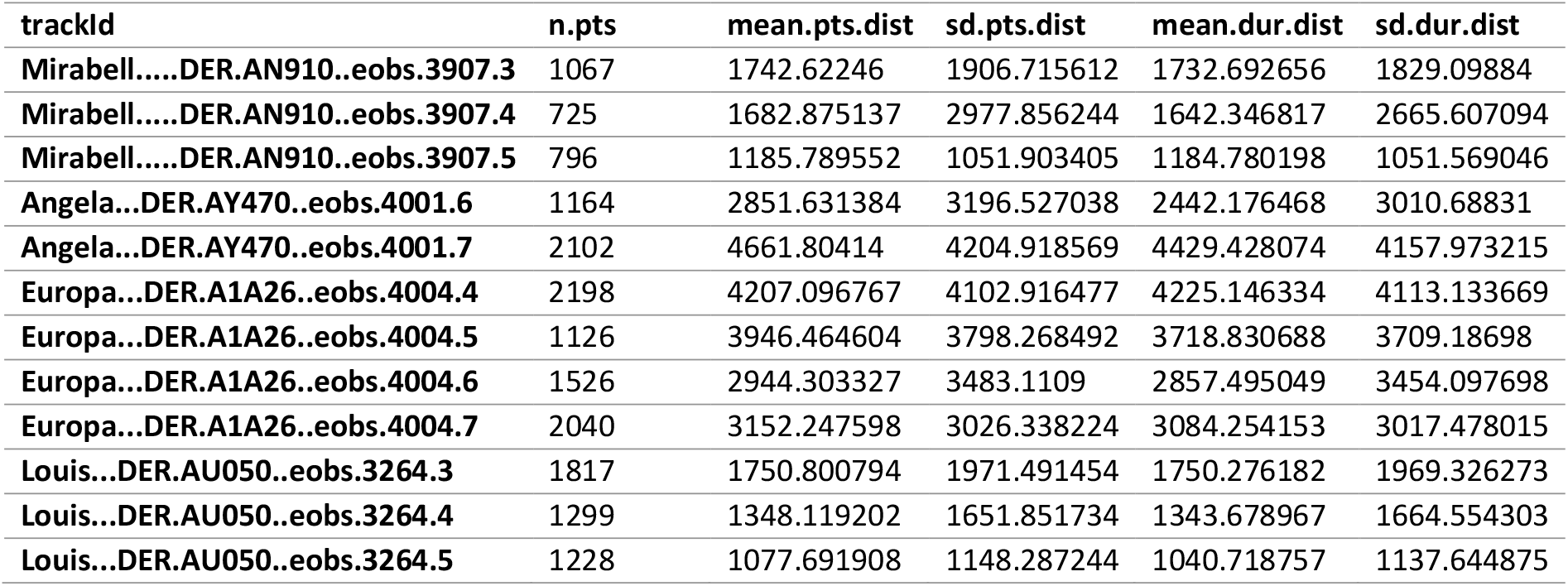
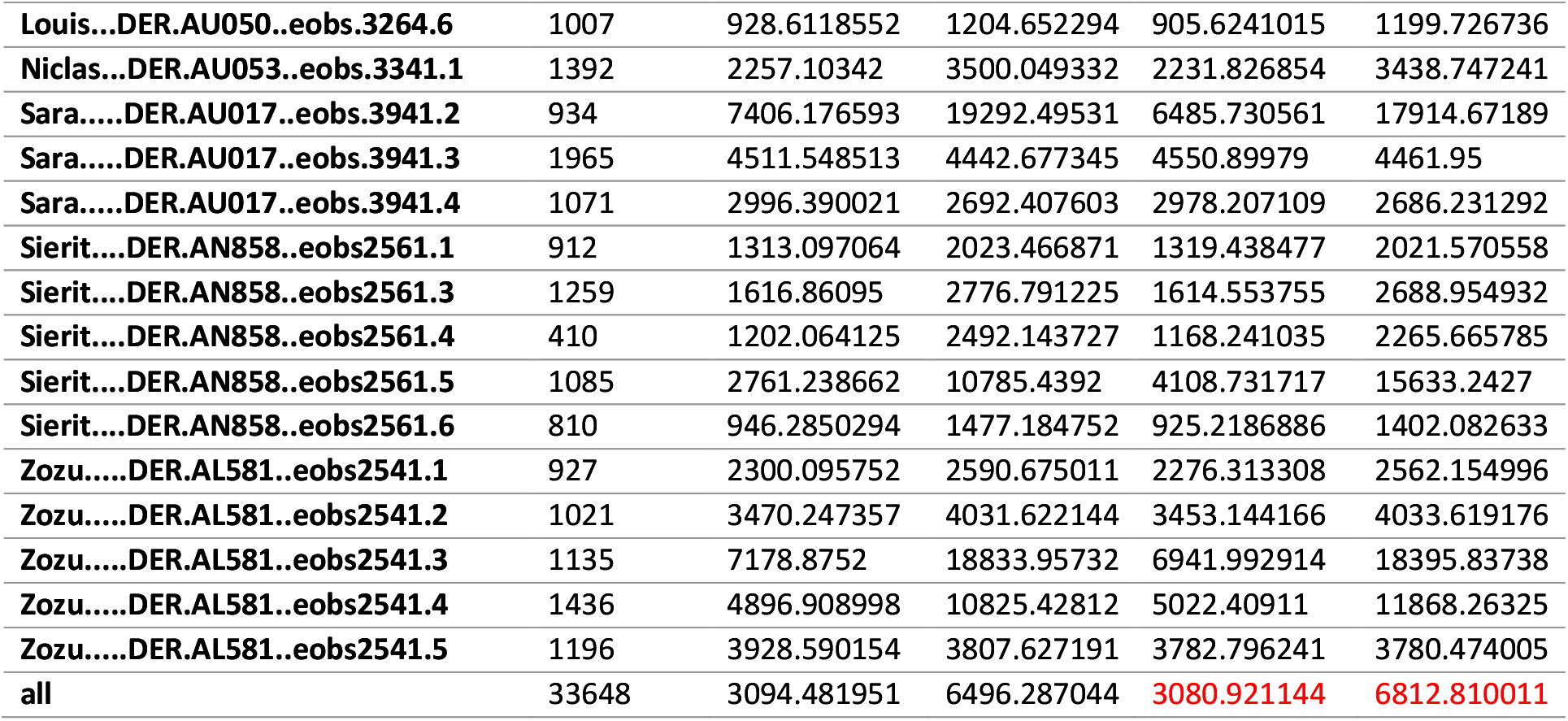
Individual distance to nest estimates (average by number of points or duration) for the selected white stork tracks per year as extracted from the Radial Area Use Around Location App.

**Table A7.** Individual distance to nest aggregates for the selected white stork tracks per year as extracted from the Radial Area Use Around Location App.

| track | radius | n.loc | prop.locs | prop.dur |
| --- | --- | --- | --- | --- |
| mean | 100 | 27 | 0.07050475 | 0.07359914 |
| mean | 200 | 27 | 0.120886264 | 0.125562264 |
| mean | 300 | 27 | 0.166305769 | 0.17283712 |
| mean | 400 | 27 | 0.209185746 | 0.21602148 |
| mean | 500 | 27 | 0.248530207 | 0.256422712 |
| mean | 600 | 27 | 0.284792939 | 0.292774072 |
| mean | 700 | 27 | 0.318143478 | 0.326442904 |
| mean | 800 | 27 | 0.349623684 | 0.358204367 |
| mean | 900 | 27 | 0.382267751 | 0.390928695 |
| mean | 1000 | 27 | 0.410282823 | 0.419254026 |
| mean | 1100 | 27 | 0.438071933 | 0.447514327 |
| mean | 1200 | 27 | 0.46430113 | 0.473516665 |
| mean | 1300 | 27 | 0.486930487 | 0.496209658 |
| mean | 1400 | 27 | 0.507347293 | 0.516780366 |
| mean | 1500 | 27 | 0.527544522 | 0.537301605 |
| mean | 1600 | 27 | 0.546703806 | 0.556488901 |
| mean | 1700 | 27 | 0.5652476 | 0.57501796 |
| mean | 1800 | 27 | 0.582880481 | 0.592778199 |
| mean | 1900 | 27 | 0.600045365 | 0.610269546 |
| mean | 2000 | 27 | 0.615787757 | 0.625899319 |
| sd | 100 | 27 | 0.038063701 | 0.037645589 |
| sd | 200 | 27 | 0.057932138 | 0.057971696 |
| sd | 300 | 27 | 0.075820615 | 0.075168051 |
| sd | 400 | 27 | 0.091722642 | 0.090976019 |
| sd | 500 | 27 | 0.104960895 | 0.104150645 |
| sd | 600 | 27 | 0.116662389 | 0.115174191 |
| sd | 700 | 27 | 0.129575757 | 0.127850287 |
| sd | 800 | 27 | 0.139312269 | 0.137399704 |
| sd | 900 | 27 | 0.14910923 | 0.14643638 |
| sd | 1000 | 27 | 0.156650479 | 0.153414763 |
| sd | 1100 | 27 | 0.162089419 | 0.157896675 |
| sd | 1200 | 27 | 0.16639207 | 0.161714689 |
| sd | 1300 | 27 | 0.170376741 | 0.165217743 |
| sd | 1400 | 27 | 0.174431474 | 0.169134056 |
| sd | 1500 | 27 | 0.177257535 | 0.171743784 |
| sd | 1600 | 27 | 0.178673537 | 0.17286471 |
| sd | 1700 | 27 | 0.180660139 | 0.174971612 |
| sd | 1800 | 27 | 0.181701215 | 0.175479897 |
| sd | 1900 | 27 | 0.184930415 | 0.178240964 |
| sd | 2000 | 27 | 0.186417296 | 0.179337544 |

## Appendix 3 individual results for red kites

**Table A8.**
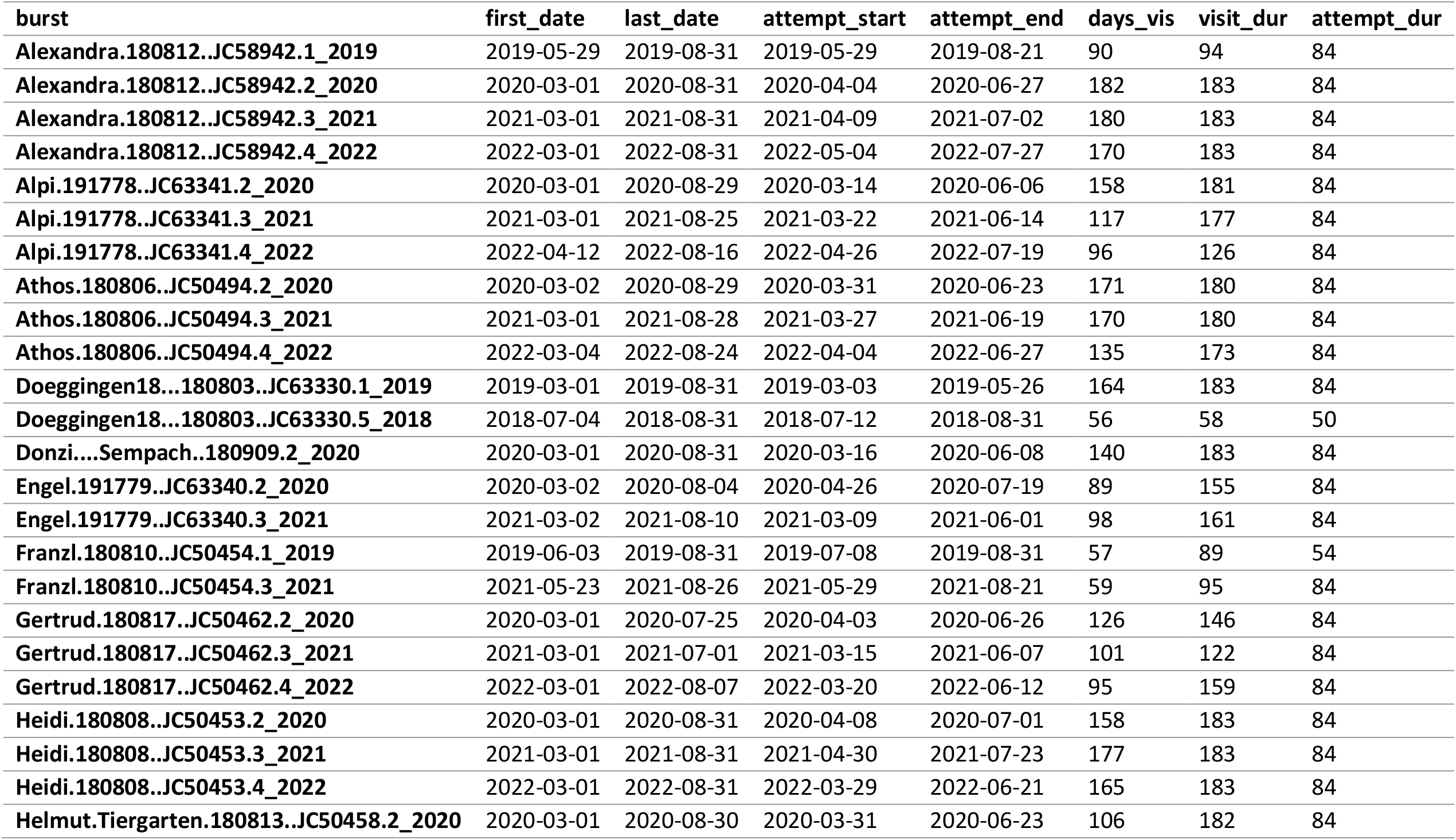
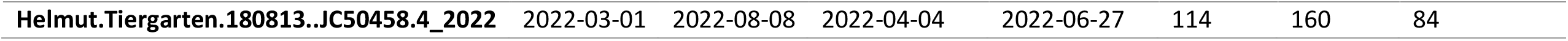
Individual nesting times and durations for the selected red kite tracks per year as extracted from the Nest Cluster Detection App.

| burst | first_date | last_date | attempt_start | attempt_end | days_vis | visit_dur | attempt_dur |
| --- | --- | --- | --- | --- | --- | --- | --- |
| Alexandra.180812..JC58942.1_2019 | 2019-05-29 | 2019-08-31 | 2019-05-29 | 2019-08-21 | 90 | 94 | 84 |
| Alexandra.180812..JC58942.2_2020 | 2020-03-01 | 2020-08-31 | 2020-04-04 | 2020-06-27 | 182 | 183 | 84 |
| Alexandra.180812..JC58942.3_2021 | 2021-03-01 | 2021-08-31 | 2021-04-09 | 2021-07-02 | 180 | 183 | 84 |
| Alexandra.180812..JC58942.4_2022 | 2022-03-01 | 2022-08-31 | 2022-05-04 | 2022-07-27 | 170 | 183 | 84 |
| Alpi.191778..JC63341.2_2020 | 2020-03-01 | 2020-08-29 | 2020-03-14 | 2020-06-06 | 158 | 181 | 84 |
| Alpi.191778..JC63341.3_2021 | 2021-03-01 | 2021-08-25 | 2021-03-22 | 2021-06-14 | 117 | 177 | 84 |
| Alpi.191778..JC63341.4_2022 | 2022-04-12 | 2022-08-16 | 2022-04-26 | 2022-07-19 | 96 | 126 | 84 |
| Athos.180806..JC50494.2_2020 | 2020-03-02 | 2020-08-29 | 2020-03-31 | 2020-06-23 | 171 | 180 | 84 |
| Athos.180806..JC50494.3_2021 | 2021-03-01 | 2021-08-28 | 2021-03-27 | 2021-06-19 | 170 | 180 | 84 |
| Athos.180806..JC50494.4_2022 | 2022-03-04 | 2022-08-24 | 2022-04-04 | 2022-06-27 | 135 | 173 | 84 |
| Doeggingen18...180803..JC63330.1_2019 | 2019-03-01 | 2019-08-31 | 2019-03-03 | 2019-05-26 | 164 | 183 | 84 |
| Doeggingen18...180803..JC63330.5_2018 | 2018-07-04 | 2018-08-31 | 2018-07-12 | 2018-08-31 | 56 | 58 | 50 |
| Donzi....Sempach..180909.2_2020 | 2020-03-01 | 2020-08-31 | 2020-03-16 | 2020-06-08 | 140 | 183 | 84 |
| Engel.191779..JC63340.2_2020 | 2020-03-02 | 2020-08-04 | 2020-04-26 | 2020-07-19 | 89 | 155 | 84 |
| Engel.191779..JC63340.3_2021 | 2021-03-02 | 2021-08-10 | 2021-03-09 | 2021-06-01 | 98 | 161 | 84 |
| Franzl.180810..JC50454.1_2019 | 2019-06-03 | 2019-08-31 | 2019-07-08 | 2019-08-31 | 57 | 89 | 54 |
| Franzl.180810..JC50454.3_2021 | 2021-05-23 | 2021-08-26 | 2021-05-29 | 2021-08-21 | 59 | 95 | 84 |
| Gertrud.180817..JC50462.2_2020 | 2020-03-01 | 2020-07-25 | 2020-04-03 | 2020-06-26 | 126 | 146 | 84 |
| Gertrud.180817..JC50462.3_2021 | 2021-03-01 | 2021-07-01 | 2021-03-15 | 2021-06-07 | 101 | 122 | 84 |
| Gertrud.180817..JC50462.4_2022 | 2022-03-01 | 2022-08-07 | 2022-03-20 | 2022-06-12 | 95 | 159 | 84 |
| Heidi.180808..JC50453.2_2020 | 2020-03-01 | 2020-08-31 | 2020-04-08 | 2020-07-01 | 158 | 183 | 84 |
| Heidi.180808..JC50453.3_2021 | 2021-03-01 | 2021-08-31 | 2021-04-30 | 2021-07-23 | 177 | 183 | 84 |
| Heidi.180808..JC50453.4_2022 | 2022-03-01 | 2022-08-31 | 2022-03-29 | 2022-06-21 | 165 | 183 | 84 |
| Helmut.Tiergarten.180813..JC50458.2_2020 | 2020-03-01 | 2020-08-30 | 2020-03-31 | 2020-06-23 | 106 | 182 | 84 |
| Helmut.Tiergarten.180813..JC50458.4_2022 | 2022-03-01 | 2022-08-08 | 2022-04-04 | 2022-06-27 | 114 | 160 | 84 |

**Table A9.** Individual flight speed estimates for the selected red kite tracks per year as extracted from the Extract Two Movement Speeds App.

| trackID | mode1 | antimode | mode2 | mean.above.antimode | sd.above.antimode |
| --- | --- | --- | --- | --- | --- |
| Alexandra.180812..JC58942.1 | 0.078416103 | 2.554415336 | 7.582833534 | 7.67753315 | 2.84198719 |
| Alexandra.180812..JC58942.2 | 0.076873708 | 2.615080172 | 6.844634114 | 7.63125178 | 2.8962374 |
| Alexandra.180812..JC58942.3 | 0.094655229 | 3.02740335 | 7.162127767 | 7.68702966 | 2.8424282 |
| Alexandra.180812..JC58942.4 | 0.096121059 | 2.280402616 | 6.968633384 | 8.13655005 | 3.13316986 |
| Alpi.191778..JC63341.1 | 0.06241733 | 1.677461749 | 6.977283931 | 7.8432528 | 3.19915325 |
| Alpi.191778..JC63341.2 | 0.053380813 | 1.509620653 | 6.810684733 | 7.77875146 | 3.33452401 |
| Alpi.191778..JC63341.3 | 0.059261584 | 2.290401121 | 7.376306781 | 7.86242063 | 3.12862823 |
| Alpi.191778..JC63341.4 | 0.069509155 | 1.630882188 | 6.993385946 | 7.83199084 | 3.2505531 |
| Athos.180806..JC50494.1 | 0.101494352 | 2.216627337 | 6.943316118 | 7.69378259 | 3.27821533 |
| Athos.180806..JC50494.2 | 0.113109496 | 1.829291928 | 5.659644484 | 7.20138304 | 3.31519538 |
| Athos.180806..JC50494.3 | 0.088763848 | 1.74728031 | 6.367846443 | 7.14497938 | 3.25656587 |
| Athos.180806..JC50494.4 | 0.085306684 | 1.734927089 | 6.162621099 | 7.27047901 | 3.29203103 |
| Doeggingen18...180803..JC63330.1 | 0.064757934 | 2.072041793 | 7.59588035 | 7.96435046 | 3.16773103 |
| Doeggingen18...180803..JC63330.5 | 0.081858775 | 2.8308902 | 7.33417003 | 8.19763904 | 2.88023945 |
| Engel.191779..JC63340.1 | 0.048295175 | 2.400119135 | 7.743161057 | 8.85218753 | 2.65099554 |
| Engel.191779..JC63340.2 | 0.053777297 | 2.415024745 | 8.447494569 | 8.97524949 | 2.63038156 |
| Engel.191779..JC63340.3 | 0.058591031 | 2.700864523 | 7.599879786 | 8.73116303 | 2.59958168 |
| Engel.191779..JC63340.4 | 0.097267412 | 2.494364604 | 7.243807399 | 8.54033222 | 2.77907006 |
| Franzl.180810..JC50454.1 | 0.108140712 | 1.941179603 | 7.058905927 | 7.71360838 | 3.26307633 |
| Franzl.180810..JC50454.2 | 0.129970398 | 2.091363736 | 6.868485031 | 7.84833854 | 3.4721482 |
| Franzl.180810..JC50454.3 | 0.121745114 | 2.46062707 | 6.971492041 | 7.93202302 | 3.48758188 |
| Franzl.180810..JC50454.4 | 0.132254218 | 1.823627622 | 6.35726251 | 7.70911307 | 3.42243451 |
| Gertrud.180817..JC50462.1 | 0.134633682 | 3.76157864 | 7.640599151 | 7.58896569 | 2.44112384 |
| Gertrud.180817..JC50462.2 | 0.120132948 | 2.79829202 | 6.208933467 | 7.48376137 | 2.9627278 |
| Gertrud.180817..JC50462.3 | 0.125235149 | 2.614983603 | 6.727840518 | 7.4082519 | 2.98038403 |
| Gertrud.180817..JC50462.4 | 0.137000753 | 2.918618299 | 6.831156265 | 8.02809292 | 3.36483611 |
| Heidi.180808..JC50453.1 | 0.095267052 | 2.057799471 | 6.949064347 | 7.28550769 | 2.9258061 |
| Heidi.180808..JC50453.2 | 0.114254859 | 1.873620854 | 6.320055704 | 7.18717918 | 3.10625246 |
| Heidi.180808..JC50453.3 | 0.122802167 | 2.229151384 | 5.9241761 | 7.29656316 | 3.10584143 |
| Heidi.180808..JC50453.4 | 0.118505282 | 1.892200889 | 6.769381999 | 7.12559636 | 2.93906782 |
| Helmut.Tiergarten.180813..JC50458.1 | 0.113228492 | 2.07116101 | 6.776659064 | 7.82314863 | 3.24284226 |
| Helmut.Tiergarten.180813..JC50458.2 | 0.092034039 | 1.927677413 | 6.506560407 | 7.51760171 | 3.44921115 |
| Helmut.Tiergarten.180813..JC50458.3 | 0.094814481 | 1.964464039 | 6.397631567 | 7.51669179 | 3.300516 |
| Helmut.Tiergarten.180813..JC50458.4 | 0.111093623 | 1.951572572 | 7.096334559 | 7.39216533 | 3.18221806 |
| Donzi....Sempach..180909.1 | 0.030600427 | 2.191926326 | 7.444370322 | 8.00238295 | 2.99605892 |
| Donzi....Sempach..180909.2 | 0.063080413 | 2.214769255 | 6.823768985 | 7.74876448 | 3.18269069 |
| mean | 0.093018078 | 2.244769796 | 6.930177486 | 7.76744673 | 3.09170849 |
| sd | 0.028197308 | 0.465977779 | 0.559864719 | 0.46014797 | 0.26453379 |

**Table A10.**
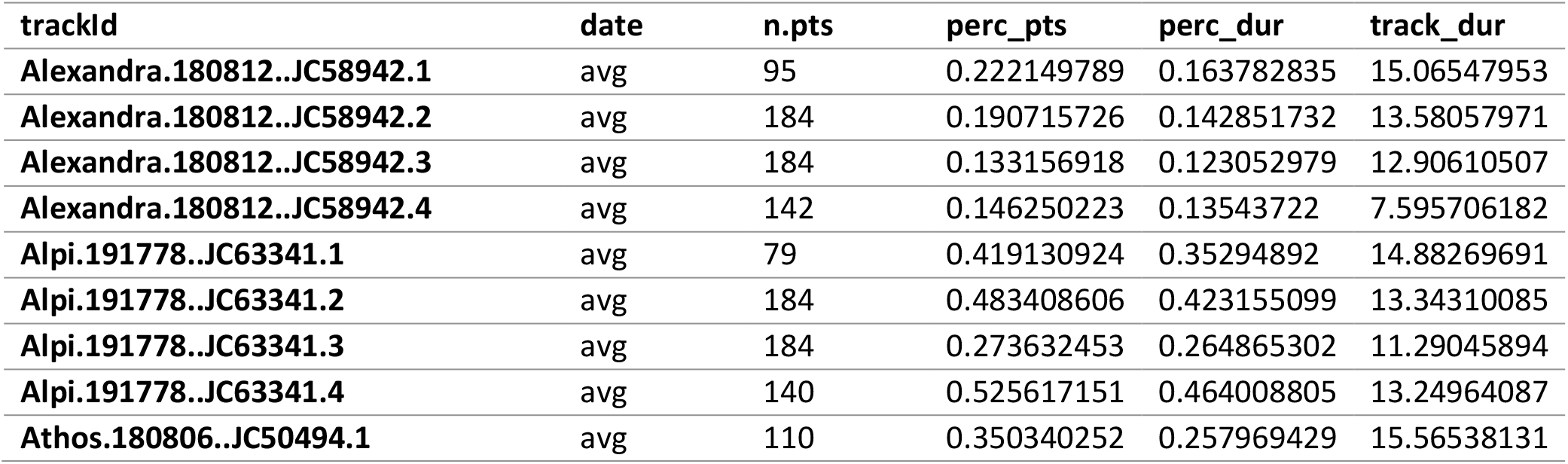
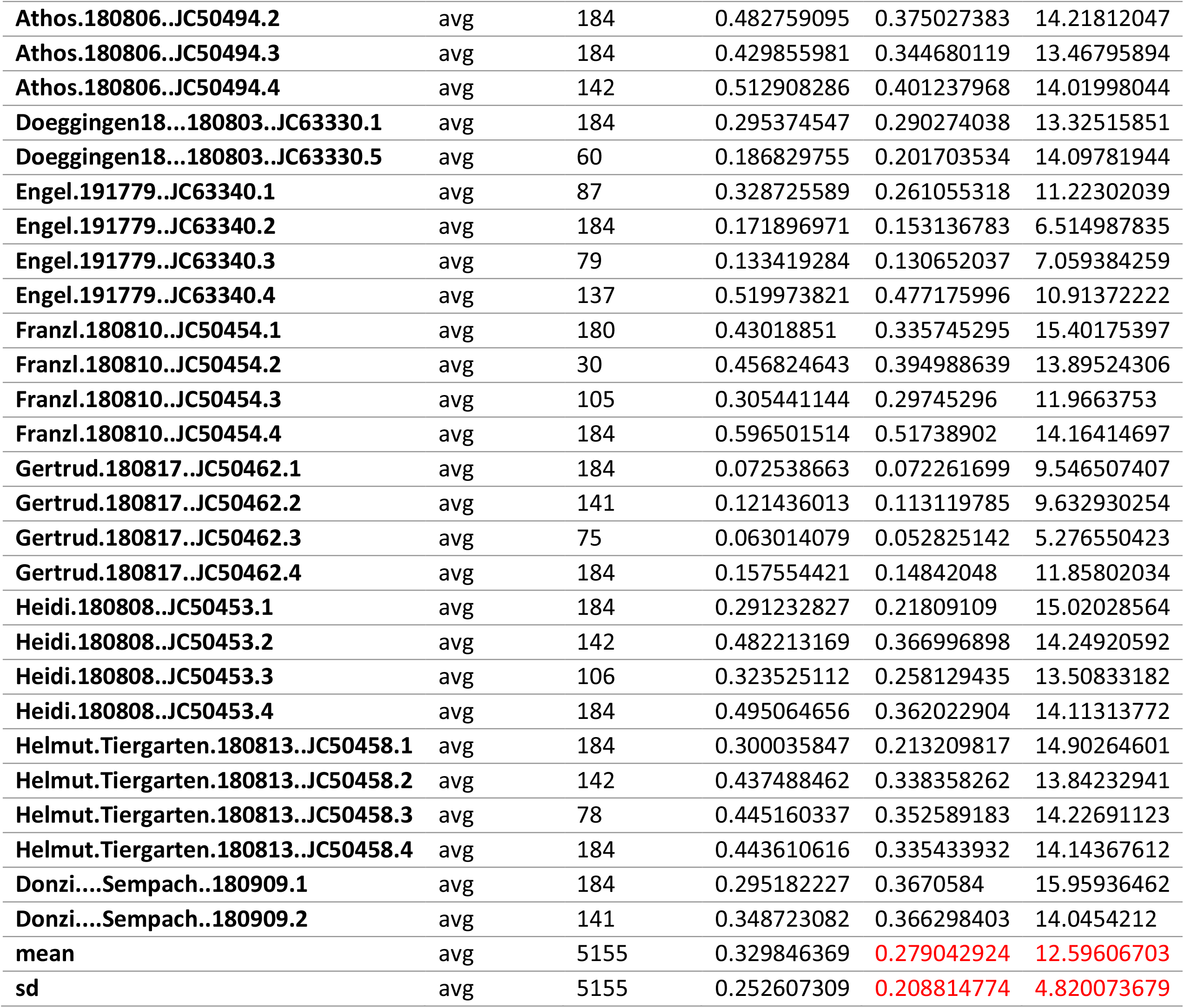
Individual daily flight duration estimates for the selected red kite tracks per year as extracted from the Daily Proportions App.

| trackId | date | n.pts | perc_pts | perc_dur | track_dur |
| --- | --- | --- | --- | --- | --- |
| Alexandra.180812..JC58942.1 | avg | 95 | 0.222149789 | 0.163782835 | 15.06547953 |
| Alexandra.180812..JC58942.2 | avg | 184 | 0.190715726 | 0.142851732 | 13.58057971 |
| Alexandra.180812..JC58942.3 | avg | 184 | 0.133156918 | 0.123052979 | 12.90610507 |
| Alexandra.180812..JC58942.4 | avg | 142 | 0.146250223 | 0.13543722 | 7.595706182 |
| Alpi.191778..JC63341.1 | avg | 79 | 0.419130924 | 0.35294892 | 14.88269691 |
| Alpi.191778..JC63341.2 | avg | 184 | 0.483408606 | 0.423155099 | 13.34310085 |
| Alpi.191778..JC63341.3 | avg | 184 | 0.273632453 | 0.264865302 | 11.29045894 |
| Alpi.191778..JC63341.4 | avg | 140 | 0.525617151 | 0.464008805 | 13.24964087 |
| Athos.180806..JC50494.1 | avg | 110 | 0.350340252 | 0.257969429 | 15.56538131 |
| Athos.180806..JC50494.2 | avg | 184 | 0.482759095 | 0.375027383 | 14.21812047 |
| Athos.180806..JC50494.3 | avg | 184 | 0.429855981 | 0.344680119 | 13.46795894 |
| Athos.180806..JC50494.4 | avg | 142 | 0.512908286 | 0.401237968 | 14.01998044 |
| Doeggingen18...180803..JC63330.1 | avg | 184 | 0.295374547 | 0.290274038 | 13.32515851 |
| Doeggingen18...180803..JC63330.5 | avg | 60 | 0.186829755 | 0.201703534 | 14.09781944 |
| Engel.191779..JC63340.1 | avg | 87 | 0.328725589 | 0.261055318 | 11.22302039 |
| Engel.191779..JC63340.2 | avg | 184 | 0.171896971 | 0.153136783 | 6.514987835 |
| Engel.191779..JC63340.3 | avg | 79 | 0.133419284 | 0.130652037 | 7.059384259 |
| Engel.191779..JC63340.4 | avg | 137 | 0.519973821 | 0.477175996 | 10.91372222 |
| Franzl.180810..JC50454.1 | avg | 180 | 0.43018851 | 0.335745295 | 15.40175397 |
| Franzl.180810..JC50454.2 | avg | 30 | 0.456824643 | 0.394988639 | 13.89524306 |
| Franzl.180810..JC50454.3 | avg | 105 | 0.305441144 | 0.29745296 | 11.9663753 |
| Franzl.180810..JC50454.4 | avg | 184 | 0.596501514 | 0.51738902 | 14.16414697 |
| Gertrud.180817..JC50462.1 | avg | 184 | 0.072538663 | 0.072261699 | 9.546507407 |
| Gertrud.180817..JC50462.2 | avg | 141 | 0.121436013 | 0.113119785 | 9.632930254 |
| Gertrud.180817..JC50462.3 | avg | 75 | 0.063014079 | 0.052825142 | 5.276550423 |
| Gertrud.180817..JC50462.4 | avg | 184 | 0.157554421 | 0.14842048 | 11.85802034 |
| Heidi.180808..JC50453.1 | avg | 184 | 0.291232827 | 0.21809109 | 15.02028564 |
| Heidi.180808..JC50453.2 | avg | 142 | 0.482213169 | 0.366996898 | 14.24920592 |
| Heidi.180808..JC50453.3 | avg | 106 | 0.323525112 | 0.258129435 | 13.50833182 |
| Heidi.180808..JC50453.4 | avg | 184 | 0.495064656 | 0.362022904 | 14.11313772 |
| Helmut.Tiergarten.180813..JC50458.1 | avg | 184 | 0.300035847 | 0.213209817 | 14.90264601 |
| Helmut.Tiergarten.180813..JC50458.2 | avg | 142 | 0.437488462 | 0.338358262 | 13.84232941 |
| Helmut.Tiergarten.180813..JC50458.3 | avg | 78 | 0.445160337 | 0.352589183 | 14.22691123 |
| Helmut.Tiergarten.180813..JC50458.4 | avg | 184 | 0.443610616 | 0.335433932 | 14.14367612 |
| Donzi....Sempach..180909.1 | avg | 184 | 0.295182227 | 0.3670584 | 15.95936462 |
| Donzi....Sempach..180909.2 | avg | 141 | 0.348723082 | 0.366298403 | 14.0454212 |
| mean | avg | 5155 | 0.329846369 | 0.279042924 | 12.59606703 |
| sd | avg | 5155 | 0.252607309 | 0.208814774 | 4.820073679 |

**Table A11.**
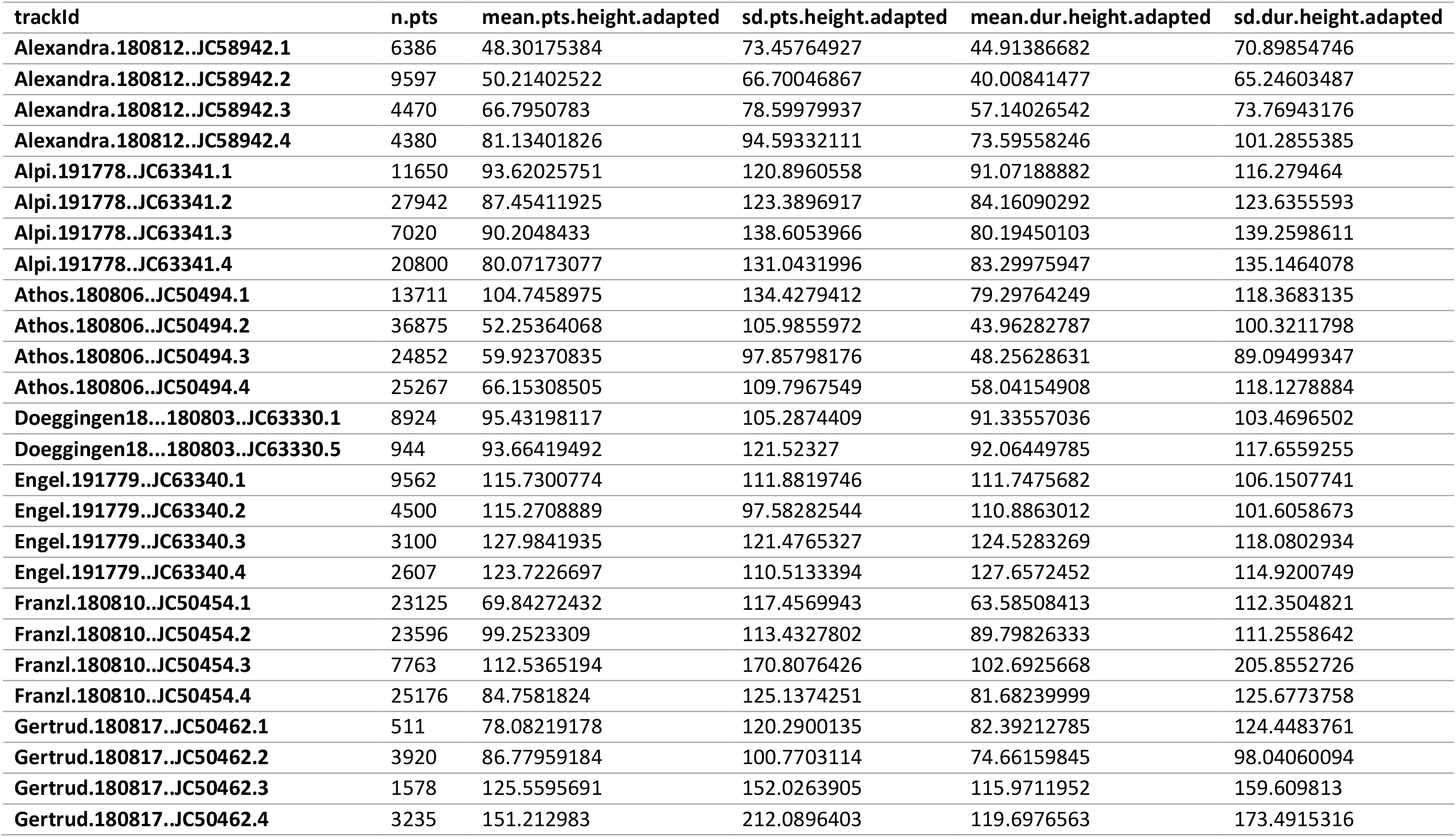
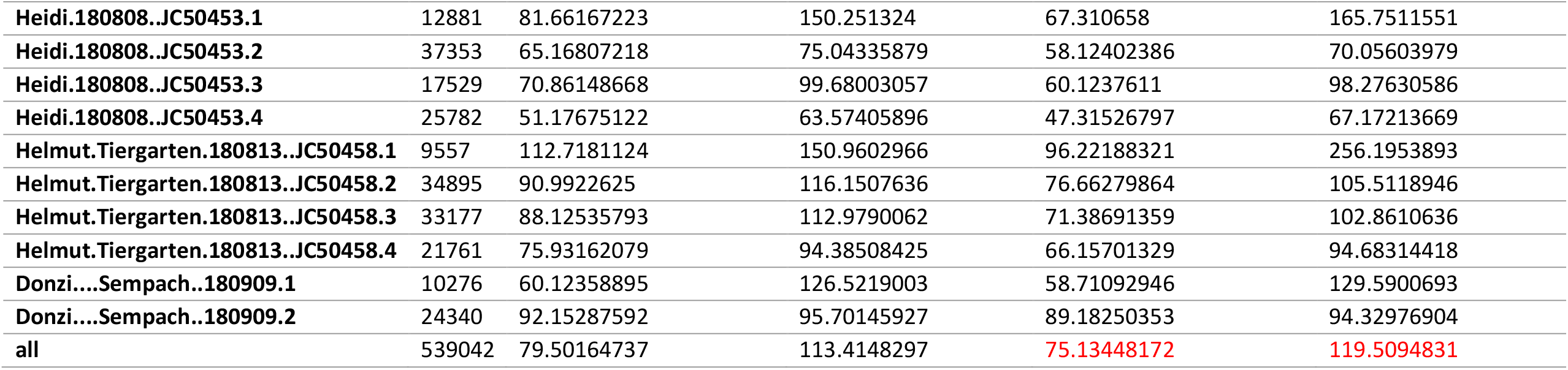
Individual flight height estimates (average by number of points or duration) for the selected red kite tracks per year as extracted from the Add Elevation and Height Above Ground App.

| trackId | n.pts | mean.pts.height.adapted | sd.pts.height.adapted | mean.dur.height.adapted | sd.dur.height.adapted |
| --- | --- | --- | --- | --- | --- |
| Alexandra.180812..JC58942.1 | 6386 | 48.30175384 | 73.45764927 | 44.91386682 | 70.89854746 |
| Alexandra.180812..JC58942.2 | 9597 | 50.21402522 | 66.70046867 | 40.00841477 | 65.24603487 |
| Alexandra.180812..JC58942.3 | 4470 | 66.7950783 | 78.59979937 | 57.14026542 | 73.76943176 |
| Alexandra.180812..JC58942.4 | 4380 | 81.13401826 | 94.59332111 | 73.59558246 | 101.2855385 |
| Alpi.191778..JC63341.1 | 11650 | 93.62025751 | 120.8960558 | 91.07188882 | 116.279464 |
| Alpi.191778..JC63341.2 | 27942 | 87.45411925 | 123.3896917 | 84.16090292 | 123.6355593 |
| Alpi.191778..JC63341.3 | 7020 | 90.2048433 | 138.6053966 | 80.19450103 | 139.2598611 |
| Alpi.191778..JC63341.4 | 20800 | 80.07173077 | 131.0431996 | 83.29975947 | 135.1464078 |
| Athos.180806..JC50494.1 | 13711 | 104.7458975 | 134.4279412 | 79.29764249 | 118.3683135 |
| Athos.180806..JC50494.2 | 36875 | 52.25364068 | 105.9855972 | 43.96282787 | 100.3211798 |
| Athos.180806..JC50494.3 | 24852 | 59.92370835 | 97.85798176 | 48.25628631 | 89.09499347 |
| Athos.180806..JC50494.4 | 25267 | 66.15308505 | 109.7967549 | 58.04154908 | 118.1278884 |
| Doeggingen18...180803..JC63330.1 | 8924 | 95.43198117 | 105.2874409 | 91.33557036 | 103.4696502 |
| Doeggingen18...180803..JC63330.5 | 944 | 93.66419492 | 121.52327 | 92.06449785 | 117.6559255 |
| Engel.191779..JC63340.1 | 9562 | 115.7300774 | 111.8819746 | 111.7475682 | 106.1507741 |
| Engel.191779..JC63340.2 | 4500 | 115.2708889 | 97.58282544 | 110.8863012 | 101.6058673 |
| Engel.191779..JC63340.3 | 3100 | 127.9841935 | 121.4765327 | 124.5283269 | 118.0802934 |
| Engel.191779..JC63340.4 | 2607 | 123.7226697 | 110.5133394 | 127.6572452 | 114.9200749 |
| Franzl.180810..JC50454.1 | 23125 | 69.84272432 | 117.4569943 | 63.58508413 | 112.3504821 |
| Franzl.180810..JC50454.2 | 23596 | 99.2523309 | 113.4327802 | 89.79826333 | 111.2558642 |
| Franzl.180810..JC50454.3 | 7763 | 112.5365194 | 170.8076426 | 102.6925668 | 205.8552726 |
| Franzl.180810..JC50454.4 | 25176 | 84.7581824 | 125.1374251 | 81.68239999 | 125.6773758 |
| Gertrud.180817..JC50462.1 | 511 | 78.08219178 | 120.2900135 | 82.39212785 | 124.4483761 |
| Gertrud.180817..JC50462.2 | 3920 | 86.77959184 | 100.7703114 | 74.66159845 | 98.04060094 |
| Gertrud.180817..JC50462.3 | 1578 | 125.5595691 | 152.0263905 | 115.9711952 | 159.609813 |
| Gertrud.180817..JC50462.4 | 3235 | 151.212983 | 212.0896403 | 119.6976563 | 173.4915316 |
| Heidi.180808..JC50453.1 | 12881 | 81.66167223 | 150.251324 | 67.310658 | 165.7511551 |
| Heidi.180808..JC50453.2 | 37353 | 65.16807218 | 75.04335879 | 58.12402386 | 70.05603979 |
| Heidi.180808..JC50453.3 | 17529 | 70.86148668 | 99.68003057 | 60.1237611 | 98.27630586 |
| Heidi.180808..JC50453.4 | 25782 | 51.17675122 | 63.57405896 | 47.31526797 | 67.17213669 |
| Helmut.Tiergarten.180813..JC50458.1 | 9557 | 112.7181124 | 150.9602966 | 96.22188321 | 256.1953893 |
| Helmut.Tiergarten.180813..JC50458.2 | 34895 | 90.9922625 | 116.1507636 | 76.66279864 | 105.5118946 |
| Helmut.Tiergarten.180813..JC50458.3 | 33177 | 88.12535793 | 112.9790062 | 71.38691359 | 102.8610636 |
| Helmut.Tiergarten.180813..JC50458.4 | 21761 | 75.93162079 | 94.38508425 | 66.15701329 | 94.68314418 |
| Donzi....Sempach..180909.1 | 10276 | 60.12358895 | 126.5219003 | 58.71092946 | 129.5900693 |
| Donzi....Sempach..180909.2 | 24340 | 92.15287592 | 95.70145927 | 89.18250353 | 94.32976904 |
| all | 539042 | 79.50164737 | 113.4148297 | 75.13448172 | 119.5094831 |

**Table A12.** Individual flight height aggregates for the selected red kite tracks per year as extracted from the Add Elevation and Height Above Ground App.

| trackId | height_threshold | n.loc | prop.locs | prop.dur |
| --- | --- | --- | --- | --- |
| mean | 25 | 36 | 0.252984645 | 0.281495426 |
| mean | 50 | 36 | 0.427408286 | 0.467569266 |
| mean | 75 | 36 | 0.568550682 | 0.60910679 |
| mean | 100 | 36 | 0.673007949 | 0.707616701 |
| mean | 125 | 36 | 0.748891351 | 0.777708394 |
| mean | 150 | 36 | 0.806954143 | 0.830627406 |
| mean | 175 | 36 | 0.851618276 | 0.870658196 |
| mean | 200 | 36 | 0.886381334 | 0.901747429 |
| mean | 225 | 36 | 0.911617942 | 0.923551144 |
| mean | 250 | 36 | 0.929796101 | 0.939794287 |
| mean | 275 | 36 | 0.94309343 | 0.950942715 |
| mean | 300 | 36 | 0.953334858 | 0.959689905 |
| sd | 25 | 36 | 0.09081007 | 0.103053612 |
| sd | 50 | 36 | 0.128718752 | 0.14444074 |
| sd | 75 | 36 | 0.138288067 | 0.146799742 |
| **sd** | 100 | 36 | 0.126684076 | 0.130872204 |
| **sd** | 125 | 36 | 0.110293827 | 0.110545679 |
| **sd** | 150 | 36 | 0.090061903 | 0.088434453 |
| **sd** | 175 | 36 | 0.070591888 | 0.067021876 |
| **sd** | 200 | 36 | 0.056116719 | 0.050729102 |
| **sd** | 225 | 36 | 0.045809701 | 0.039359787 |
| **sd** | 250 | 36 | 0.039989217 | 0.032337125 |
| **sd** | 275 | 36 | 0.033500839 | 0.027385352 |
| **sd** | 300 | 36 | 0.029147108 | 0.023791069 |

**Table A13.** Individual distance to nest estimates (average by number of points or duration) for the selected red kite tracks per year as extracted from the Radial Area Use Around Location App.

| **trackId** | **n.pts** | **mean.pts.dist** | **sd.pts.dist** | **mean.dur.dist** | **sd.dur.dist** |
| --- | --- | --- | --- | --- | --- |
| **Alexandra.180812..JC58942.1** | 6386 | 751.0614826 | 623.6066781 | 703.2024518 | 631.4281449 |
| **Alexandra.180812..JC58942.2** | 9597 | 862.1619183 | 1298.238258 | 796.9390955 | 1580.993373 |
| **Alexandra.180812..JC58942.3** | 4470 | 1373.730624 | 1955.626174 | 1139.63976 | 1629.287561 |
| **Alexandra.180812..JC58942.4** | 4380 | 2153.843866 | 2225.517043 | 1853.119079 | 2447.761472 |
| **Alpi.191778..JC63341.2** | 27942 | 2238.118676 | 1480.904826 | 2127.017284 | 1552.031704 |
| **Alpi.191778..JC63341.3** | 7020 | 2039.619934 | 1432.606126 | 2024.98084 | 1510.262327 |
| **Alpi.191778..JC63341.4** | 20800 | 2857.689472 | 1857.657157 | 2849.700194 | 2134.603322 |
| **Athos.180806..JC50494.2** | 36875 | 1408.527385 | 1280.893796 | 1273.034215 | 1213.601687 |
| **Athos.180806..JC50494.3** | 24852 | 1066.496583 | 1114.266566 | 958.5543989 | 1010.487934 |
| **Athos.180806..JC50494.4** | 25267 | 1451.800344 | 1307.032032 | 1338.710031 | 1250.420278 |
| **Doeggingen18...180803..JC63330.1** | 8924 | 1691.822776 | 1693.005901 | 1345.216386 | 1419.070841 |
| **Doeggingen18...180803..JC63330.5** | 944 | 2069.460785 | 1996.489768 | 1867.599185 | 1913.643624 |
| **Engel.191779..JC63340.2** | 4500 | 2830.459531 | 2065.645676 | 2324.245936 | 1867.769282 |
| **Engel.191779..JC63340.3** | 3100 | 2561.217111 | 1781.560379 | 2409.355805 | 1818.58691 |
| **Franzl.180810..JC50454.1** | 23125 | 2017.713809 | 1286.678825 | 1904.421491 | 1267.122161 |
| **Franzl.180810..JC50454.3** | 7763 | 1114.373765 | 854.271747 | 1072.322515 | 781.4718712 |
| Gertrud.180817..JC50462.2 | 3920 | 2530.442065 | 3483.385401 | 2146.385989 | 3593.504313 |
| Gertrud.180817..JC50462.3 | 1578 | 3957.86202 | 6385.197036 | 5064.315347 | 8340.596222 |
| Gertrud.180817..JC50462.4 | 3235 | 15963.11641 | 23737.41356 | 10093.82803 | 19636.44954 |
| Heidi.180808..JC50453.2 | 37353 | 1023.601409 | 725.2710695 | 965.0148633 | 680.4352799 |
| Heidi.180808..JC50453.3 | 17529 | 823.4765648 | 821.5943227 | 762.9543466 | 680.3933399 |
| Heidi.180808..JC50453.4 | 25782 | 937.802497 | 631.9091974 | 876.7770308 | 620.7226158 |
| Helmut.Tiergarten.180813..JC50458.2 | 34895 | 1708.835836 | 2087.400152 | 1446.245039 | 1807.59872 |
| Helmut.Tiergarten.180813..JC50458.4 | 21761 | 1131.819677 | 1827.382515 | 960.4486328 | 1460.565866 |
| Donzi....Sempach..180909.2 | 24340 | 1280.508748 | 1541.571685 | 1263.026192 | 1498.734168 |
| all | 386338 | 1638.943214 | 3011.134749 | 1648.236639 | 3425.905407 |

**Table A14.** Individual distance to nest aggregates for the selected red kite tracks per year as extracted from the Radial Area Use Around Location App.

| track | radius | n.loc | prop.locs | prop.dur |
| --- | --- | --- | --- | --- |
| mean | 100 | 25 | 0.040843017 | 0.055892435 |
| mean | 200 | 25 | 0.081346655 | 0.10385573 |
| mean | 300 | 25 | 0.123055746 | 0.150652603 |
| mean | 400 | 25 | 0.167344464 | 0.198959502 |
| mean | 500 | 25 | 0.21663106 | 0.252538977 |
| mean | 600 | 25 | 0.266628759 | 0.307302176 |
| mean | 700 | 25 | 0.316701842 | 0.358624595 |
| mean | 800 | 25 | 0.366409489 | 0.410976378 |
| mean | 900 | 25 | 0.415694982 | 0.46266233 |
| mean | 1000 | 25 | 0.461647255 | 0.510328262 |
| mean | 1100 | 25 | 0.505850878 | 0.554361528 |
| mean | 1200 | 25 | 0.547037587 | 0.595001088 |
| mean | 1300 | 25 | 0.579806986 | 0.627492508 |
| mean | 1400 | 25 | 0.607563063 | 0.654187041 |
| mean | 1500 | 25 | 0.631384566 | 0.677115352 |
| mean | 1600 | 25 | 0.652937381 | 0.698378359 |
| mean | 1700 | 25 | 0.673103977 | 0.717984083 |
| mean | 1800 | 25 | 0.691844899 | 0.735786055 |
| mean | 1900 | 25 | 0.708738414 | 0.751586261 |
| mean | 2000 | 25 | 0.723896534 | 0.765383878 |
| sd | 100 | 25 | 0.027837202 | 0.036856338 |
| sd | 200 | 25 | 0.049594735 | 0.059932761 |
| sd | 300 | 25 | 0.067123395 | 0.076336964 |
| sd | 400 | 25 | 0.082967872 | 0.090466731 |
| sd | 500 | 25 | 0.099549654 | 0.106487459 |
| sd | 600 | 25 | 0.116006598 | 0.12136159 |
| sd | 700 | 25 | 0.131042802 | 0.135401164 |
| sd | 800 | 25 | 0.147175081 | 0.150008496 |
| sd | 900 | 25 | 0.162236117 | 0.164101785 |
| sd | 1000 | 25 | 0.176140409 | 0.176745213 |
| sd | 1100 | 25 | 0.18815893 | 0.186175952 |
| sd | 1200 | 25 | 0.200315892 | 0.194833528 |
| sd | 1300 | 25 | 0.208842007 | 0.200427541 |
| sd | 1400 | 25 | 0.212689629 | 0.201262814 |
| sd | 1500 | 25 | 0.214734926 | 0.201147272 |
| sd | 1600 | 25 | 0.214421575 | 0.199320644 |
| sd | 1700 | 25 | 0.21287395 | 0.196422324 |
| sd | 1800 | 25 | 0.210708014 | 0.19269737 |
| sd | 1900 | 25 | 0.208297277 | 0.188904529 |
| sd | 2000 | 25 | 0.204784965 | 0.184721834 |

## Appendix 4 Individual results for marsh harriers

**Table A15.** Individual nesting times and durations for the selected marsh harrier tracks per year as extracted from the Nest Cluster Detection App.

| burst | first_date | last_date | attempt_start | attempt_end | days_vis | visit_dur | attempt_dur |
| --- | --- | --- | --- | --- | --- | --- | --- |
| Arnhem.3806.6962_2021 | 2021-04-30 | 2021-06-22 | 2021-05-15 | 2021-06-22 | 44 | 53 | 38 |
| L1778022_2021 | 2021-03-27 | 2021-08-13 | 2021-04-03 | 2021-06-11 | 113 | 139 | 69 |
| L1434515_2014 | 2014-04-23 | 2014-08-20 | 2014-06-08 | 2014-08-16 | 87 | 119 | 69 |
| X53256676_2015 | 2015-04-16 | 2015-08-25 | 2015-04-16 | 2015-06-24 | 71 | 131 | 69 |
| X53256677_2016 | 2016-04-04 | 2016-08-25 | 2016-04-05 | 2016-06-13 | 91 | 143 | 69 |
| X53256678_2017 | 2017-03-28 | 2017-08-18 | 2017-03-31 | 2017-06-08 | 115 | 143 | 69 |
| X53256679_2018 | 2018-04-01 | 2018-07-10 | 2018-04-01 | 2018-06-09 | 99 | 100 | 69 |
| X54464654_2013 | 2013-04-09 | 2013-08-24 | 2013-05-17 | 2013-07-25 | 75 | 137 | 69 |
| X54464655_2014 | 2014-04-12 | 2014-08-09 | 2014-04-14 | 2014-06-22 | 92 | 119 | 69 |
| X54464656_2015 | 2015-04-09 | 2015-08-30 | 2015-04-09 | 2015-06-17 | 112 | 143 | 69 |
| X54464657_2016 | 2016-04-05 | 2016-08-10 | 2016-04-05 | 2016-06-13 | 103 | 127 | 69 |
| X544646510_2012 | 2012-05-10 | 2012-08-20 | 2012-05-10 | 2012-07-18 | 67 | 102 | 69 |

**Table A16.** Individual flight speed estimates for the selected marsh harrier tracks per year as extracted from the Extract Two Movement Speeds App. The light blue lettering indicates tracks with too little data for proper estimation. They were not used for averages.

| trackID | mode1 | antimode | mode2 | mean.above.antimode | sd.above.antimode |
| --- | --- | --- | --- | --- | --- |
| Arnhem.3806.6962 | 0.015221417 | 1.451123428 | 4.595434401 | 5.68321982 | 2.42095137 |
| L1778022 | 0.084860695 | 2.166817365 | 4.97554631 | 6.28793498 | 2.99077672 |
| L1434515 | 0.24443836 | 2.804995077 | 3.748086097 | 6.35577235 | 2.6451877 |
| X53256676 | 0.837185787 | 28.89568597 | 33.02904508 | 33.1066152 | 1.09443323 |
| X53256677 | 0.846157339 | 25.84026433 | 28.24706631 | 29.6334069 | 2.56059637 |
| X53256678 | 0.320877362 | 3.545982525 | 4.229384518 | 7.21108536 | 2.83592211 |
| X53256679 | 0.252260478 | 2.951248926 | 5.404250138 | 7.15120415 | 2.85497083 |
| **X54464654** | 0.230009119 | 2.487456281 | 3.917736897 | 6.27879953 | 2.82224069 |
| **X54464655** | 0.219455598 | 2.151055093 | 4.072785101 | 5.80620708 | 2.71258104 |
| **X54464656** | 0.21889318 | 1.944921307 | 3.251798683 | 5.71936407 | 2.91416815 |
| **X54464657** | 0.225016018 | 1.92859362 | 3.803864372 | 5.79383563 | 2.75086869 |
| **X544646510** | 0.146292822 | 1.49823355 | 3.492233086 | 5.49273314 | 2.75975484 |
| **mean** | 0.303389015 | 6.472198122 | 8.563935916 | 10.3766815 | 2.61353765 |
| **sd** | 0.264079413 | 9.800379561 | 10.37879808 | 9.84903367 | 0.50295589 |
| **Mean adapted** | 0.195732505 | 2.293042717 | 4.14911196 |  |  |
| **Sd adapted** | 0.084664995 | 0.627187381 | 0.635614019 |  |  |

**Table A17.** Individual daily flight duration estimates for the selected marsh harrier tracks per year as extracted from the Daily Proportions App.

| trackId | date | n.pts | perc_pts | perc_dur | track_dur |
| --- | --- | --- | --- | --- | --- |
| **Arnhem.3806.6962** | avg | 39 | 0.339251799 | 0.340352799 | 17.44940171 |
| **L1778022** | avg | 69 | 0.325740686 | 0.322055331 | 16.98095411 |
| **L1434515** | avg | 69 | 0.286494569 | 0.265656135 | 22.92785427 |
| **X53256676** | avg | 70 | 0.294783695 | 0.290318898 | 14.08550794 |
| **X53256677** | avg | 70 | 0.38658365 | 0.368959698 | 14.91770635 |
| **X53256678** | avg | 70 | 0.38908549 | 0.281514526 | 22.46332143 |
| **X53256679** | avg | 70 | 0.386860221 | 0.268976699 | 23.40265873 |
| **X54464654** | avg | 70 | 0.326073163 | 0.354607535 | 13.6872471 |
| **X54464655** | avg | 67 | 0.381733253 | 0.422749199 | 13.53555159 |
| **X54464656** | avg | 70 | 0.389976716 | 0.422029737 | 12.81669569 |
| **X54464657** | avg | 67 | 0.446450464 | 0.471627981 | 11.59402576 |
| **X544646510** | avg | 69 | 0.382209296 | 0.401751639 | 13.1665 |
| **mean** | avg | 800 | 0.362179211 | 0.350559689 | 16.39996701 |
| **sd** | avg | 800 | 0.137931349 | 0.136445065 | 5.02393791 |

**Table A18.**
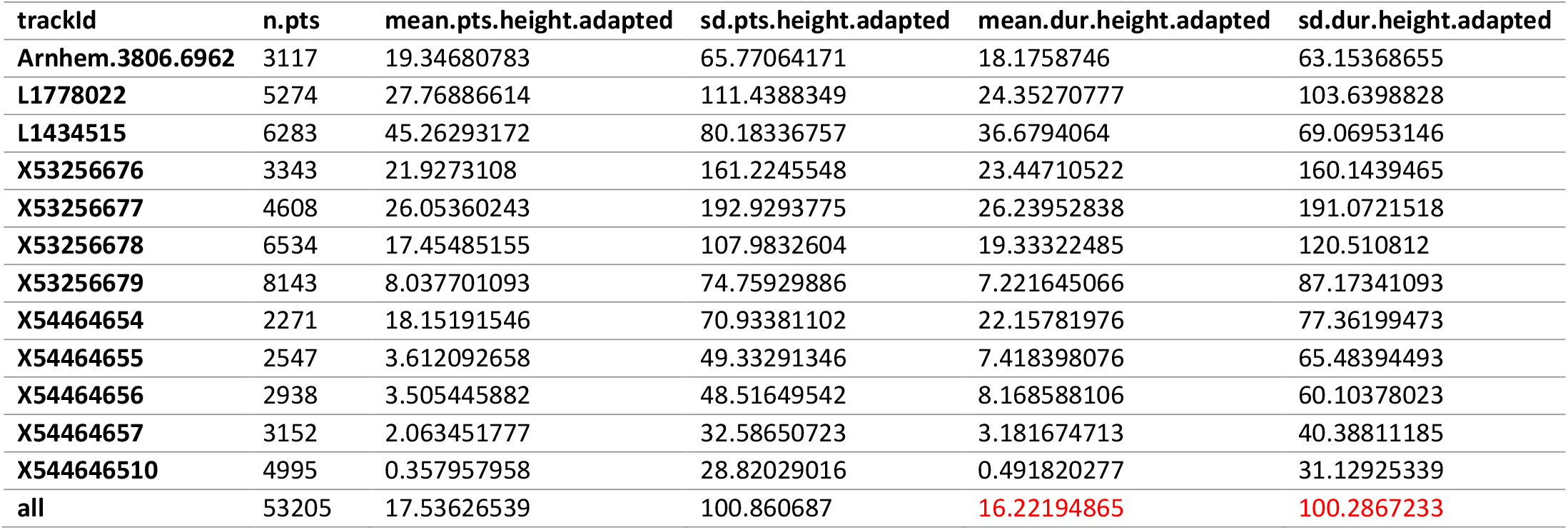
Individual flight height estimates (average by number of points or duration) for the selected marsh harrier tracks per year as extracted from the Add Elevation and Height Above Ground App.

**Table A19.**
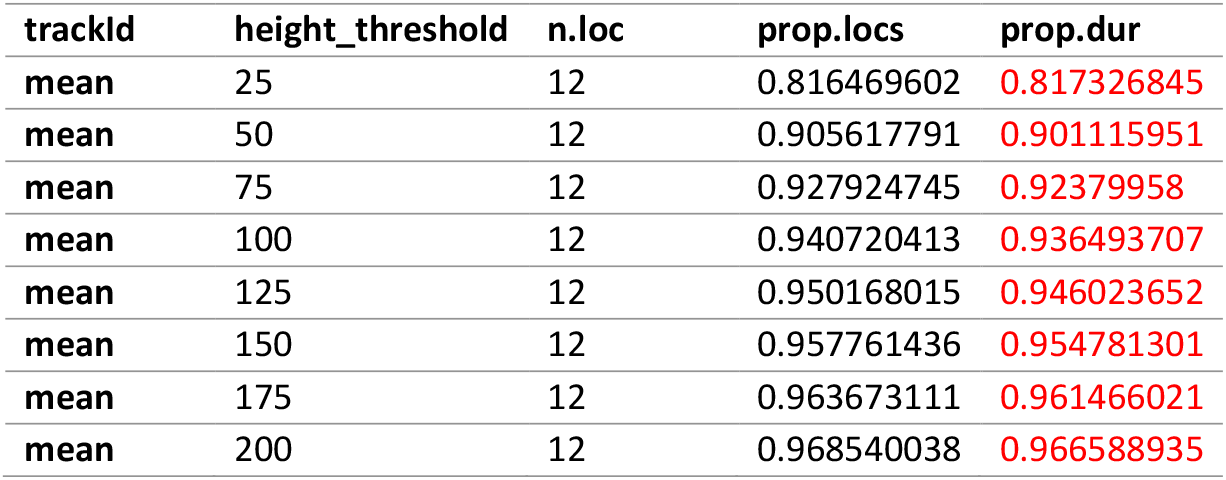
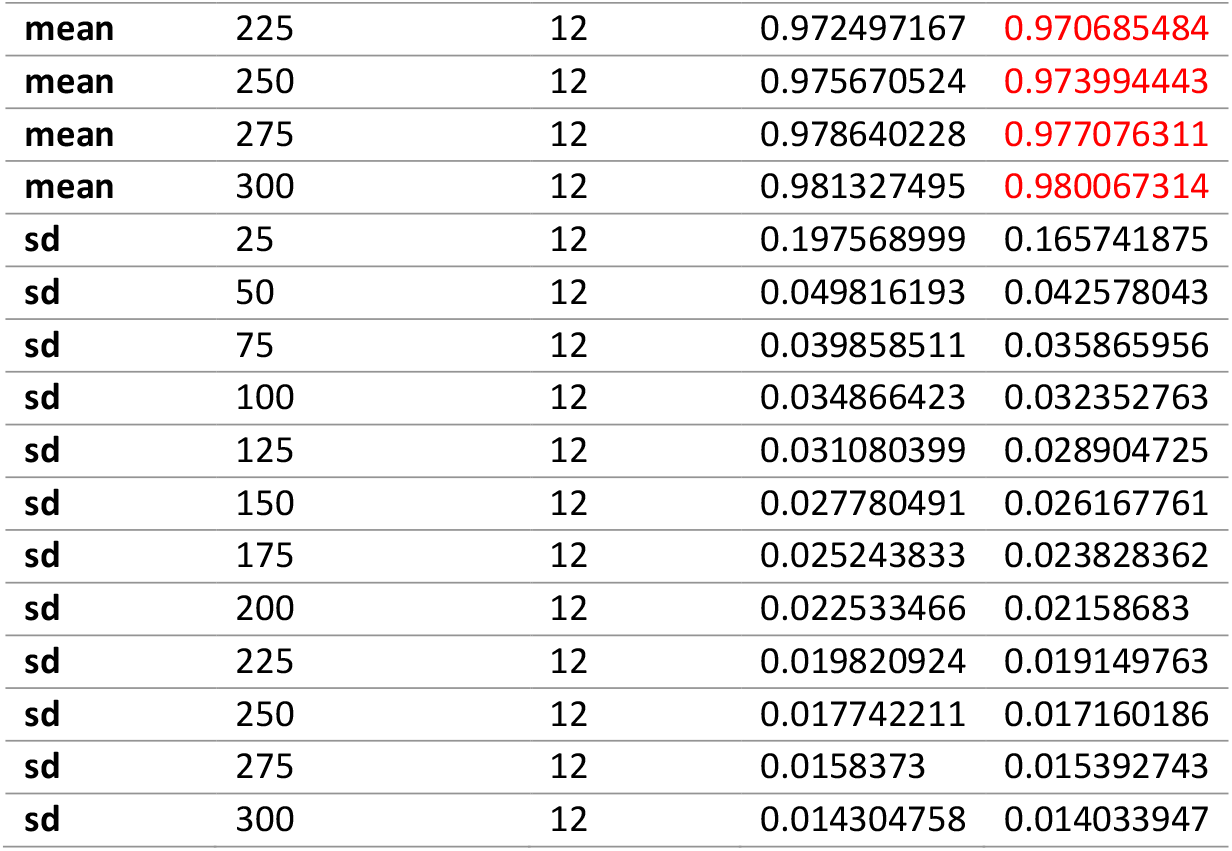
Individual flight height aggregates for the selected marsh harrier tracks per year as extracted from the Add Elevation and Height Above Ground App.

| trackId | height_threshold | n.loc | prop.locs | prop.dur |
| --- | --- | --- | --- | --- |
| mean | 25 | 12 | 0.816469602 | 0.817326845 |
| mean | 50 | 12 | 0.905617791 | 0.901115951 |
| mean | 75 | 12 | 0.927924745 | 0.92379958 |
| mean | 100 | 12 | 0.940720413 | 0.936493707 |
| mean | 125 | 12 | 0.950168015 | 0.946023652 |
| mean | 150 | 12 | 0.957761436 | 0.954781301 |
| mean | 175 | 12 | 0.963673111 | 0.961466021 |
| mean | 200 | 12 | 0.968540038 | 0.966588935 |
| mean | 225 | 12 | 0.972497167 | 0.970685484 |
| mean | 250 | 12 | 0.975670524 | 0.973994443 |
| mean | 275 | 12 | 0.978640228 | 0.977076311 |
| mean | 300 | 12 | 0.981327495 | 0.980067314 |
| sd | 25 | 12 | 0.197568999 | 0.165741875 |
| sd | 50 | 12 | 0.049816193 | 0.042578043 |
| sd | 75 | 12 | 0.039858511 | 0.035865956 |
| sd | 100 | 12 | 0.034866423 | 0.032352763 |
| sd | 125 | 12 | 0.031080399 | 0.028904725 |
| sd | 150 | 12 | 0.027780491 | 0.026167761 |
| sd | 175 | 12 | 0.025243833 | 0.023828362 |
| sd | 200 | 12 | 0.022533466 | 0.02158683 |
| sd | 225 | 12 | 0.019820924 | 0.019149763 |
| sd | 250 | 12 | 0.017742211 | 0.017160186 |
| sd | 275 | 12 | 0.0158373 | 0.015392743 |
| sd | 300 | 12 | 0.014304758 | 0.014033947 |

**Table A20.** Individual distance to nest estimates (average by number of points or duration) for the selected marsh harrier tracks per year as extracted from the Radial Area Use Around Location App.

| trackId | n.pts | mean.pts.dist | sd.pts.dist | mean.dur.dist | sd.dur.dist |
| --- | --- | --- | --- | --- | --- |
| Arnhem.3806.6962 | 3117 | 2044.238455 | 2246.815171 | 2022.814105 | 2295.028707 |
| L1778022 | 5274 | 1223.196737 | 1661.241723 | 1211.950855 | 1627.808984 |
| L1434515 | 6283 | 802.793675 | 1190.075852 | 673.4729556 | 829.4674018 |
| X53256676 | 3343 | 1067.848768 | 1155.542806 | 1066.925193 | 1155.368746 |
| X53256677 | 4608 | 566.6210552 | 820.107474 | 558.7637513 | 817.5874984 |
| X53256678 | 6534 | 957.4518128 | 1149.14105 | 894.9059715 | 1131.220994 |
| X53256679 | 8143 | 1039.2242 | 927.3842445 | 989.1938806 | 1087.973637 |
| X54464654 | 2271 | 1254.408488 | 1056.969603 | 1236.915763 | 1091.979924 |
| X54464655 | 2547 | 883.6642834 | 863.4222424 | 926.3313248 | 960.2971028 |
| **X54464656** | 2938 | 1267.473988 | 3537.774295 | 1640.438258 | 5920.671432 |
| **X54464657** | 3152 | 1228.520948 | 1012.013318 | 1270.686568 | 1072.260654 |
| **X544646510** | 4995 | 1267.249794 | 1840.997683 | 1220.49261 | 1756.561316 |
| **all** | 53205 | 1086.207847 | 1557.392534 | **1100.219297** | **2121.767743** |

**Table A21.** Individual distance to nest aggregates for the selected marsh harrier tracks per year as extracted from the Radial Area Use Around Location App.

| **track** | **radius** | **n.loc** | **prop.locs** | **prop.dur** |
| --- | --- | --- | --- | --- |
| **mean** | 100 | 12 | 0.117905537 | **0.133620161** |
| **mean** | 200 | 12 | 0.209134822 | **0.225810655** |
| **mean** | 300 | 12 | 0.284788721 | **0.301196095** |
| **mean** | 400 | 12 | 0.348504893 | **0.365276685** |
| **mean** | 500 | 12 | 0.409904292 | **0.423676567** |
| **mean** | 600 | 12 | 0.461010026 | **0.471369152** |
| **mean** | 700 | 12 | 0.509415152 | **0.516928298** |
| **mean** | 800 | 12 | 0.549409807 | **0.554316347** |
| **mean** | 900 | 12 | 0.583213275 | **0.586758172** |
| **mean** | 1000 | 12 | 0.618955971 | **0.622017194** |
| **mean** | 1100 | 12 | 0.655574825 | **0.660393066** |
| **mean** | 1200 | 12 | 0.686934483 | **0.692328728** |
| **mean** | 1300 | 12 | 0.715433377 | **0.721793015** |
| **mean** | 1400 | 12 | 0.73996643 | **0.745920346** |
| **mean** | 1500 | 12 | 0.759782319 | **0.764501627** |
| **mean** | 1600 | 12 | 0.778657831 | **0.781897479** |
| **mean** | 1700 | 12 | 0.796440578 | **0.79937824** |
| **mean** | 1800 | 12 | 0.81441656 | **0.81623592** |
| **mean** | 1900 | 12 | 0.831499624 | **0.832171942** |
| **mean** | 2000 | 12 | 0.845781419 | **0.845473887** |
| **sd** | 100 | 12 | 0.055117435 | 0.055737719 |
| **sd** | 200 | 12 | 0.088477212 | 0.088005713 |
| **sd** | 300 | 12 | 0.118631442 | 0.115787831 |
| **sd** | 400 | 12 | 0.131995941 | 0.130253822 |
| **sd** | 500 | 12 | 0.137823738 | 0.136419793 |
| **sd** | 600 | 12 | 0.138431856 | 0.135497112 |
| **sd** | 700 | 12 | 0.139957864 | 0.13560008 |
| **sd** | 800 | 12 | 0.145590737 | 0.14117296 |
| **sd** | 900 | 12 | 0.148427765 | 0.142022731 |
| **sd** | 1000 | 12 | 0.14267115 | 0.135528323 |
| **sd** | 1100 | 12 | 0.137354307 | 0.134961064 |
| **sd** | 1200 | 12 | 0.133033017 | 0.132549317 |
| **sd** | 1300 | 12 | 0.127365938 | 0.127975187 |
| **sd** | 1400 | 12 | 0.120569651 | 0.121741135 |
| **sd** | 1500 | 12 | 0.113927537 | 0.115979575 |
| **sd** | 1600 | 12 | 0.106818823 | 0.109902853 |
| **sd** | 1700 | 12 | 0.101405187 | 0.104603781 |
| **sd** | 1800 | 12 | 0.095571821 | 0.098275751 |
| **sd** | 1900 | 12 | 0.090923448 | 0.092411079 |
| **sd** | 2000 | 12 | 0.088141765 | 0.088952223 |

